# Precursor-Dependent Routing of Aromatic Amino Acids Determines Lignin Structure in Grasses by Sensitivity-Enhanced Solid-State NMR

**DOI:** 10.64898/2026.01.17.700116

**Authors:** Priya Sahu, Debkumar Debnath, Peng Xiao, Shubha S. Gunaga, Faith J. Scott, Max Bentelspacher, Yifan Xu, Frederic Mentink-Vigier, Jaime Barros, Tuo Wang

## Abstract

Lignin biosynthesis in grasses exhibits unique metabolic flexibility, yet the precursor-specific routing of carbon into lignin polymers remains poorly resolved *in planta*. Here, we combine ^13^C-isotope labeling with solid-state NMR under sensitivity-enhancement by dynamic nuclear polarization (DNP), to directly track phenylalanine- and tyrosine-derived carbon incorporation into the lignin polymer in *Brachypodium distachyon*. Precursor-specific ^13^C labeling reveals that phenylalanine is the dominant contributor to canonical guaiacyl and syringyl lignins, whereas tyrosine preferentially enriches hydroxyphenyl lignin and hydroxycinnamates, including ferulates characteristic of grass cell walls. Two-dimensional ^13^C-^13^C correlation NMR resolves distinct lignin moieties arising from each precursor. Disruption of *p*-coumarate 3-hydroxylase (C3H) selectively impairs phenylalanine-derived lignification, while tyrosine-derived lignin remains comparatively unchanged, maintaining polymer assembly through alternative metabolic routes. These findings show precursor-dependent control of lignin composition and reveal tyrosine-mediated lignification as a compensatory pathway in grasses. This work also establishes precursor-resolved solid-state NMR and DNP as a powerful framework for dissecting lignin biosynthesis and metabolic plasticity in plant cell walls.

**SIGNIFICANCE STATEMENT:** Lignin is a complex plant polymer that strengthens cell walls but also limits the efficiency of biomass processing for agriculture and bioenergy. Grasses possess a unique lignin biosynthetic flexibility that is not well understood. By combining stable isotope labeling with solid-state NMR spectroscopy, we directly traced how the aromatic amino acids, phenylalanine and tyrosine, contribute differently to lignin formation in intact grass cell walls. We show that phenylalanine primarily builds conventional lignin structures, whereas tyrosine supplies alternative phenolic components and maintains lignin synthesis even when a key biosynthetic enzyme is disrupted. This metabolic flexibility helps explain the unique structural aspects of grass cell walls and identifies precursor-level control as a promising strategy for engineering lignin composition to improve biomass utilization.

## INTRODUCTION

Lignin is one of the most structurally complex and abundant biopolymers in terrestrial plant biomass, second only to cellulose (1, 2). Located in the secondary cell walls of vascular plants, it provides mechanical support, hydrophobicity for water transport, and defense against pathogens (3). These functions are essential for plant viability and productivity in food and bioenergy crops, and reflect the evolutionary advantage conferred by lignification during land plant colonization (4, 5). At the same time, lignin’s heterogeneous and highly cross-linked polyphenolic structure presents a major barrier to biomass conversion and utilization (6, 7). In the context of biofuel production, its resistance to enzymatic saccharification and chemical extraction limits the efficient release of fermentable sugars, posing challenges for sustainable bioenergy and bioproduct development (8, 9).

At the biochemical level, lignin is synthesized through the phenylpropanoid pathway, which converts the aromatic amino acids phenylalanine (Phe) and, in monocots, also tyrosine (Tyr), into monolignol precursors. These intermediates, including p-coumaryl alcohol, coniferyl alcohol, and sinapyl alcohol, give rise to the hydroxyphenyl (H), guaiacyl (G), and syringyl (S) monolignol subunits, respectively (10–12). After synthesis, monolignols are exported to the cell wall and oxidatively polymerized through radical-mediated coupling reactions to form the lignin macromolecule (12–14). The pathway comprises sequential deamination, hydroxylation, and methylation steps catalyzed by a suite of specialized enzymes, including phenylalanine ammonia-lyase (PAL), cinnamate 4-hydroxylase (C4H), *p*-coumarate 3-hydroxylase (C3H), *p*4-coumarate-CoA ligase (4CL), caffeoyl shikimate esterase (CSE), ferulate 5-hydroxylase (F5H), and caffeate O-methyltransferase (COMT) (12).

In dicotyledonous plants such as *Arabidopsis thaliana,* flux into the phenylpropanoid pathway occurs exclusively through Phe via PAL (15–17). Interestingly, commelinid monocots, including grasses, have bifunctional phenylalanine/tyrosine ammonia-lyases (PTALs), which can deaminate both phenylalanine and tyrosine to generate cinnamic acid and *p*-coumaric acid, respectively (18–21). This additional entry point introduces greater metabolic flexibility and may contribute to the distinct structural and compositional features of grass lignins, including their elevated proportion of hydroxyphenyl subunits and the unique incorporation of tricin and other flavonoid-derived moieties (22–24). Despite these observations, the relative contributions of Phe-and Tyr-derived precursors to lignin biosynthesis in grasses remain poorly resolved. In particular, it is unclear to what extent Phe or Tyr-derived *p*-coumarate pools are diverted into the monolignol pathway versus diverted the flavonoid or soluble phenolic branches, and how metabolic flux through these dual entry points responds to genetic perturbations (9, 25). These uncertainties limit efforts to rationally engineer lignin composition for improved biomass digestibility and processing efficiency (26, 27).

Stable isotope labeling offers a powerful approach to trace metabolic flux and to distinguish the fates of individual precursor pools (17, 28, 29). Feeding ^13^C-labeled Phe or ^13^C-Tyr enables monitoring of label incorporation into lignin and downstream phenylpropanoid metabolites using mass spectrometry (18, 30, 31). To extend these molecular insights to intact cell walls, complementary approaches are required that can probe the native polymeric architecture of lignin, where molecular packing, interunit linkages, and interactions with polysaccharides are preserved.

To achieve this, here we use solid-state NMR spectroscopy, which allows direct probing of insoluble and heterogeneous biopolymers *in situ* without chemical extraction or degradation (32–34). Solid-state NMR has been widely used to investigate the organization of cellulose, hemicellulose, and lignin in plant cell walls and to characterize interactions between polysaccharides and lignin (35–38). Nevertheless, the intrinsically low sensitivity of solid-state NMR, particularly for ^13^C nuclei, has constrained its application in isotope-tracing studies, especially when labeling levels are low or spectral complexity is high. Therefore, we combined solid-state NMR with Dynamic Nuclear Polarization (DNP), a sensitivity-enhancement technique that transfers polarization from unpaired electrons to NMR-active nuclei under microwave irradiation, increasing signal intensity by one to two orders of magnitude in complex biological solids (39–44). When coupled with 2D correlation experiments, DNP-enhanced solid-state NMR has been employed to enable detailed mapping of incorporation sites, interunit linkages, and spatial organization within lignocellulose (45–48). To date, most solid-state NMR studies have focused on static architecture rather than how metabolic inputs shape polymer structure in planta. The present study bridges this gap and directly links metabolic entry points to lignin polymer composition within intact biomass.

This integrated approach is applied to *Brachypodium distachyon*, a model grass that uses the most common C3 form of photosynthesis and is widely used for bioenergy and developmental research due to its small genome, short life cycle, and phylogenetic proximity to major cereals (49). In addition to wild-type plants, we examine a C3H mutant that disrupts a key 3-hydroxylation step in the phenylpropanoid pathway. C3H catalyzes the conversion of p-coumaric acid to caffeic acid, a precursor leading to G and S monolignols; loss-of-function mutations reduce lignin content and shift composition, making this mutant an informative system to track precursor incorporation (50). By feeding ^13^C-labeled Phe and Tyr to *Brachypodium* plants, we track the incorporation of both precursors into lignin in both genotypes. This work provides new insight into the differential utilization of aromatic amino acid precursors in grass lignification and into how pathway perturbation reshapes flux through the phenylpropanoid pathway. It also establishes a method combining isotopic labeling, reverse genetic analysis, and DNP-enhanced solid-state NMR to understand lignin biosynthesis during cell wall assembly and to support future efforts in plant engineering for improved energy and biomaterial applications.

## RESULTS

### Precursor-specific ^13^C labeling reveals tissue-dependent lignin incorporation

To investigate how aromatic amino acid precursors contribute to lignin biosynthesis, we used isotope labeling strategies using uniformly ^13^C-labeled phenylalanine, tyrosine, and glucose in *Brachypodium distachyon*. Samples were labeled individually with each precursor, as well as with a combination of all three to separate general metabolic carbon incorporation from lignin-specific biosynthetic flux. We first evaluated the impact of ^13^C-glucose labeling as a reference for bulk metabolism (51). As expected, the resulting ^13^C CP-MAS spectrum was dominated by intense signals in the 60-105 ppm region, corresponding to widespread incorporation of glucose-derived carbon into cellulose and hemicellulose (**Figure 1A**). In contrast, lignin-derived aromatic signals were comparatively weak and only became apparent upon magnification of the 110-190 ppm region, where resonances characteristic of G, ferulate (FA), and S units were observed (**Figure 1B**). The addition of ^13^C-Phe and ^13^C-Tyr did not alter the carbohydrate region but produced discernible changes in aromatic signal intensity and pattern (**Figure 1B**), indicating that aromatic amino acids contribute specifically to lignin labeling beyond bulk glucose metabolism.

**Figure 1.**
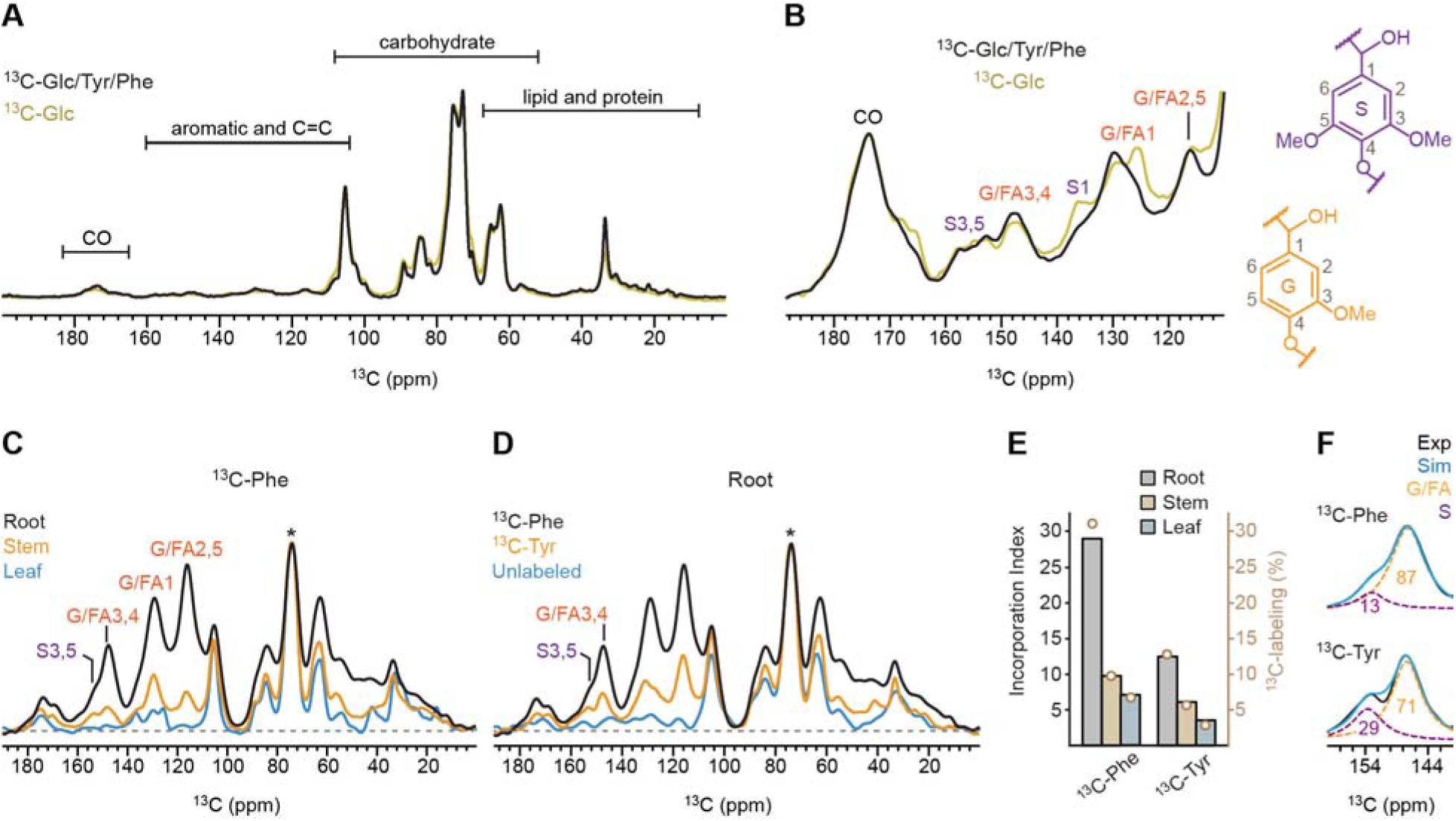
^13^C Solid-state NMR reveals differential aromatic precursor incorporation into lignin. (**A**) 1D ^13^C CP spectra of *Brachypodium* root samples labeled with ^13^C-Phe, ^13^C-Tyr, and ^13^C-glucose (black), or with only ^13^C-glucose (pale yellow). (**B**) Magnified view of the aromatic and carbonyl region showing carbon signals of guaiacyl (G), ferulate (FA), and syringyl (S) monolignol units. The simplified structures and carbon numbers of G and S are shown on the right side. (**C**) 1D ^13^C CP spectra of ^13^C-Phe-labeled root (black), stem (orange), and leaf (blue). The asterisk denotes the dominant carbohydrate peak at 73 ppm that is not ^13^C-enriched (from natural abundance of ^13^C present in unlabeled carbohydrates); this peak was used as the reference for intensity normalization. (**D**) Overlay of 1D ^13^C CP spectra collected on root samples enriched using ^13^C-Phe (black) and ^13^C-Tyr (orange), alongside an unlabeled control (blue). Labeling with ^13^C-Phe yields higher aromatic signal intensity than ^13^C-Tyr. (**E**) Histogram of lignin incorporation index values reflecting relative extent of ^13^C enrichment in lignin in each sample relative to unlabeled sample, calculated from aromatic-region integrals (108-165 ppm) normalized to the unlabeled control and scaled to the 73-ppm carbohydrate peak (the approach detailed in Methods section). The open circles in camel represent the ^13^C-labeling percentage of each sample, projected to the y-axis on the right (Supplementary Text). (**F**) Spectral deconvolution of the partially overlapping G/FA3,4 (orange dashed) and S3,5 (purple dashed) peaks in ^13^C-Phe-labeled (top) and ^13^C-Tyr-labeled (bottom) root samples. Numbers indicate the molar percentages of S and G units. Experimental (Exp; black) and simulated (Sim; blue) spectra are shown.

The aromatic amino acids, phenylalanine and tyrosine, enter lignin biosynthesis through the phenylpropanoid pathway (31). In grasses, the presence of PTAL allows both amino acids to be directly deaminated and routed toward monolignol formation (18, 20) This metabolic feature provides an opportunity to compare how different aromatic amino acid precursors are incorporated into lignin and to examine how carbons from Phe and Tyr are distributed across plant tissues (52).

We therefore examined tissue-specific incorporation in root, stem, and leaf tissues. The 1D ^13^C CP spectra revealed distinct aromatic resonances near 153 ppm, 130 ppm, and 115 ppm, corresponding to G/FA- and S-lignin carbons (**Figure 1C**). After normalization to the unlabeled carbohydrate resonance, root tissues exhibited the highest aromatic signal intensity, followed by stem and then leaf. This is consistent with established lignification gradients in grasses, where vascular bundles and pericycle tissues in roots undergo extensive secondary cell wall deposition and lignin biosynthesis (24, 53–55).

A parallel analysis of ^13^C-Tyr-labeled samples revealed the same tissue-dependent trend, with roots showing the highest aromatic signal intensity and substantially weaker signals in stems and leaves (**Figure S1**). This trend was consistently observed across all tissues examined, with aromatic signal intensity decreasing from root to stem to leaf, which confirmed that lignification is tightly regulated at the tissue level, independent of precursor identity. Given the robust aromatic labeling in roots, we focused subsequent analyses on Phe- and Tyr-labeled root tissues.

Direct comparison with an unlabeled control confirmed that both ^13^C-Phe and ^13^C-Tyr enhanced lignin-associated signals, as evidenced by the appearance of resonances assigned to S and G/FA units (**Figure 1D**). Although grasses are generally known to contain a higher proportion of S units than G units, S-lignin peaks observed in our spectra were weak across all tissues compared with G/FA, suggesting that phenylalanine and tyrosine are preferentially incorporated into G rather than S units. Notably, Phe-labeled roots consistently exhibited higher aromatic signal intensity than Tyr-labeled roots across lignin-specific regions, indicating more efficient routing of phenylalanine-derived carbon into lignin. This observation is consistent with the higher phenylalanine ammonia-lyase activity and greater flux through phenylalanine in grass phenylpropanoid metabolism (20, 26, 31). In contrast, the lower aromatic signal intensity observed with tyrosine labeling may reflect differences in precursor uptake, PTAL kinetics, or diversion of tyrosine into competing metabolic pathways (56, 57).

To assess precursor utilization, we defined a lignin incorporation index representing the fold increase in aromatic lignin signal relative to the unlabeled control, normalized to the highest carbohydrate peak. After accounting for natural ^13^C abundance and residual unlabeled lignin contributions, this index was converted to ^13^C-labeling percentages (**Figure 1E**; **Supplementary Text**). In root tissues, the lignin incorporation index decreased from 29 to 12 when the precursor was switched from ^13^C-Phe to ^13^C-Tyr, corresponding to a reduction in labeling from 31% to 13%. Thus, phenylalanine exhibits approximately 2-3-fold higher incorporation into lignin than tyrosine. A similar reduction in precursor incorporation was observed in stems and leaves (**Figure S2**).

Beyond differences in overall incorporation efficiency, the two precursors also showed distinct preferences for lignin subunit incorporation. Spectral deconvolution of the aromatic region revealed changes in the relative contributions of signals in G/FA carbons 3 and 4 at 147 ppm versus S-lignin carbons 3 and 5 at 153 ppm. In ^13^C-Phe-labeled roots, G/FA units accounted for 87% of this region, whereas S units comprised 13% (**Figure 1F**). In contrast, ^13^C-Tyr-labeled roots showed a shifted ratio of 71:29. These differences suggest precursor-dependent modulation of lignin composition, with ^13^C-Tyr being incorporated more efficiently than ^13^C-Phe into S units.

### DNP-enhanced 2D correlation spectra reveal precursor-specific lignin substructures

The limited sample quantity and partial ^13^C-labeling precluded multidimensional correlation analysis using conventional solid-state NMR. Therefore, we employed DNP to overcome this sensitivity limitation, which enhances NMR intensity by transferring polarization from unpaired electrons in the stable biradical Asympol-POK to nearby nuclei (39, 40, 58, 59). The DNP enhancement, quantified by comparing spectra acquired with and without microwave irradiation, was approximately 11-fold for the ^13^C-Phe-labeled root sample (**Figure 2A**) and 8-fold for the ^13^C-Tyr-labeled root, with enhancement factors of 38-42 observed for other *Brachypodium* samples (**Figure S3**). This reduces NMR experimental times by 100-1600-fold (**Figure S4**), thereby enabling collection of 2D ^13^C-^13^C correlation spectra for detailed analysis of lignin structure.

**Figure 2.**
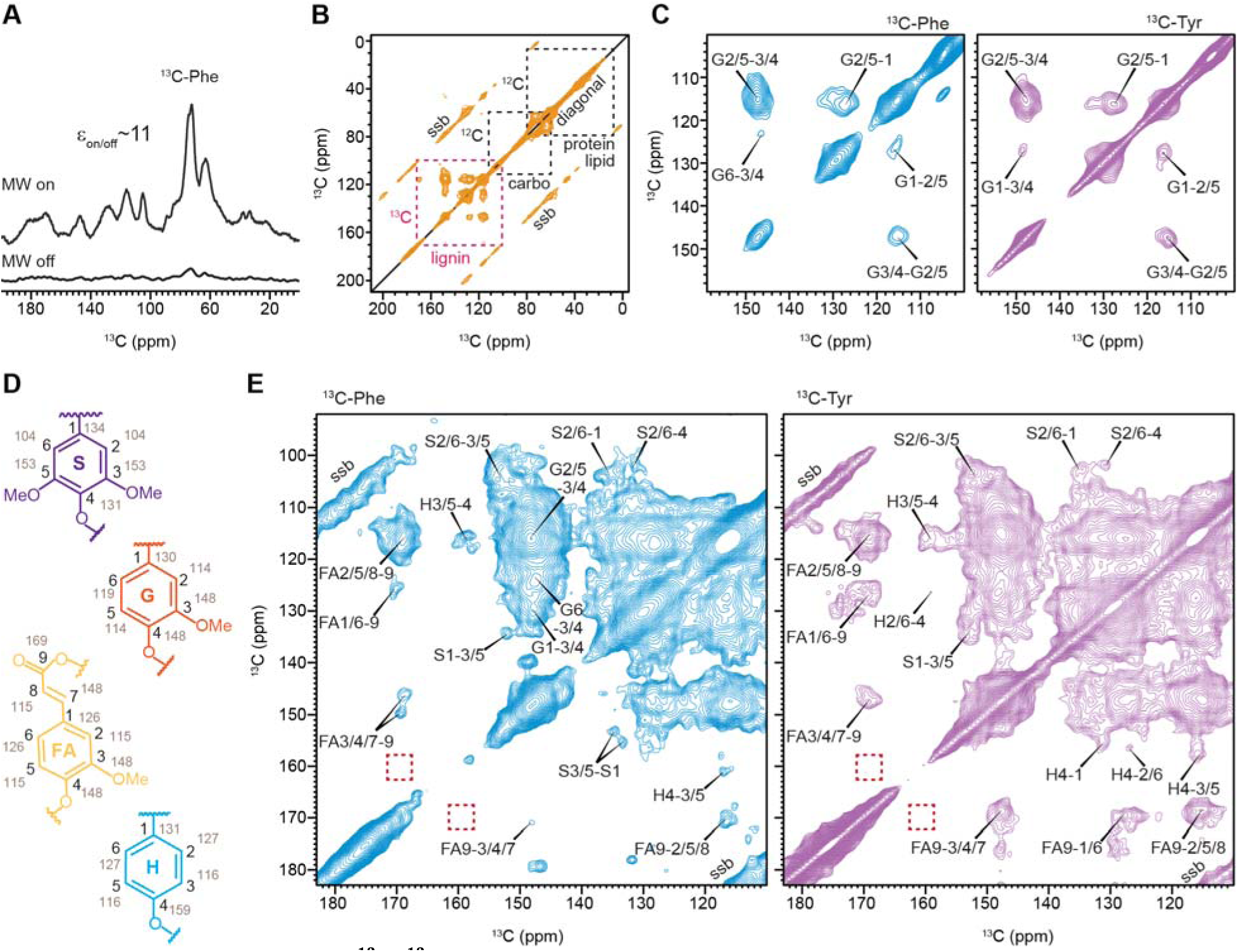
DNP-enabled 2D ^13^C-^13^C spectra reveals precursor-specific lignin labeling in wildtype root. (**A**) Dynamic nuclear polarization (DNP) enhancement of NMR sensitivity in 1D ^13^C CP spectra of ^13^C-Phe-labeled wild-type roots acquired with (top) and without (bottom) microwave (MW) irradiation. The DNP enhancement factor (ε_on/off_) is 11. (**B**) Representative full 2D ^13^C-^13^C correlation spectrum measured with a 100 ms DARR mixing time on a ^13^C-Tyr-labeled wildtype root sample. Black dashed boxes highlight unlabeled (^12^C) protein/lipid and carbohydrate signals arising from natural isotopic abundance, which show only diagonal peaks and their spinning sidebands (ssb). The magenta dashed box highlights ^13^C-labeled lignin, which exhibits additional off-diagonal intramolecular cross-peaks. (**C**) High-contour plots of the lignin region from 2D spectra of wildtype root samples labeled with ^13^C-Phe (blue) or ^13^C-Tyr (purple). The predominant signals correspond to G-lignin. (**D**) Simplified structures of different monolignol units, with carbon numbering in black and corresponding ^13^C chemical shifts annotated in grey. (**E**) Low-contour plots of the lignin region from 2D spectra of wild-type root samples labeled with ^13^C-Phe (blue) or ^13^C-Tyr (purple), revealing additional signals from S, FA, and H units. Red dashed boxes indicate characteristic positions expected for *p*CA units, which were not observed, likely due to their low abundance.

The resulting 2D ^13^C-^13^C correlation spectra exhibited intense intramolecular cross-peaks arising from lignin, due to effective incorporation of ^13^C-labeled precursors. In contrast, carbohydrates, proteins, and lipids remained largely unlabeled and therefore contributed primarily to diagonal signals and their associated spinning sidebands, without generating off-diagonal cross-peaks (**Figure 2B**). Therefore, these labeling schemes allowed for unambiguous resolution of lignin structure against the complex cellular background.

At low contour thresholds, which emphasize dominant molecules while filtering out minor components, the spectra were dominated by signals characteristic of G-lignin (**Figure 2C**). Prominent cross-peaks included correlations between C2/5 and C1 (G2/5-1) and between the C2/5 and C3/4 (G2/5-3/4). These features were consistently observed in root samples labeled with either ^13^C-phenylalanine or ^13^C-tyrosine, demonstrating that both precursors contribute to G-lignin incorporation in *Brachypodium* cell walls.

As the contour threshold was elevated for detection of lower-abundance components, numerous additional cross-peaks emerged in both root samples, corresponding to S-lignin, H-lignin, and FA units (**Figure 2D, E**). Ferulate-derived signals were readily identified by correlations involving its carbonyl carbon (C9), which resonates at 168-170 ppm, including FA2/5/8-9, FA1/6-9, and FA3/4/7-9 cross-peaks (13, 60). Signals from syringyl lignin were resolved primarily through its distinctive aromatic carbons C2 and C6, which resonate uniquely upfield at 102-108 ppm, as well as through C3 and C5, the methoxylated ring carbons resonating at 153-155 ppm. Accordingly, the characteristic S2/6-3/5 correlation appeared at approximately (104 ppm, 153 ppm), accompanied by additional resolved cross-peaks such as S2/6-1 and S2/6-4. Hydroxyphenyl lignin was identified by H3/5-4 and H2/6-4 cross-peaks, resolved through its C4 carbon at around 160 ppm. This pronounced change in chemical shift arises from strong electron withdrawal (deshielding) of the aromatic carbon directly bonded to a phenolic oxygen, further accentuated by the absence of methoxy substitution, which leaves C4 more electronically exposed.

Compared to ^13^C-Phe-labeled root, the ^13^C-Tyr-labeled root exhibited relatively stronger FA signals (**Figure 2E**), indicating that tyrosine preferentially labels ferulate moieties, which are abundant constituents of grass cell walls (61–63). We next searched for signals from *p*-coumarate (*p*CA), a closely related hydroxycinnamate that lacks the ring methoxy substitution present in FA. However, we did not observe cross-peaks at the characteristic *p*CA C4/C9 chemical shifts (160 ppm, 168 ppm; dashed boxes in **Figure 2E**) (64). Because these carbons are only three-bond apart, such correlations should be detectable if *p*CA were abundant; therefore, the absence of the C4-C9 cross peak suggests that the fraction or labeling percentage of *p*CA was low under these conditions. While plants grown in soil or pots in the greenhouse have shown a reasonable level of *p*CA, the lack of their signals here could be due to the growth stage used here, namely five-week-old plantlets, or to the growing conditions using culture tubes.

### Analysis of C3H mutant reveals precursor-specific sensitivity to lignin pathway disruption

To determine how disruption of the phenylpropanoid pathway alters precursor-specific lignin incorporation, we examined ^13^C-phenylalanine- and ^13^C-tyrosine-labeling in C3H-knockdown mutant of *Brachypodium* (50). C3H catalyzes the 3-hydroxylation of *p*-coumaric to caffeic acid, a central step required for the biosynthesis of guaiacyl and syringyl monolignols (31, 50). In root tissues, comparison with wild-type plants revealed a pronounced reduction in aromatic signal intensity in the C3H-deficient mutant for ^13^C-phenylalanine-labeled samples (**Figure 3A**), indicating substantially diminished incorporation of phenylalanine-derived carbon into lignin when C3H activity is disrupted. In contrast, this reduction was not observed in ^13^C-tyrosine-labeled roots, which showed only subtle differences in aromatic signal intensity between wild-type and C3H genotypes. This divergence suggests that tyrosine-derived lignin biosynthesis is comparatively insensitive to C3H loss, potentially reflecting its distinct metabolic routing or redirection toward H-lignin (52, 65).

**Figure 3.**
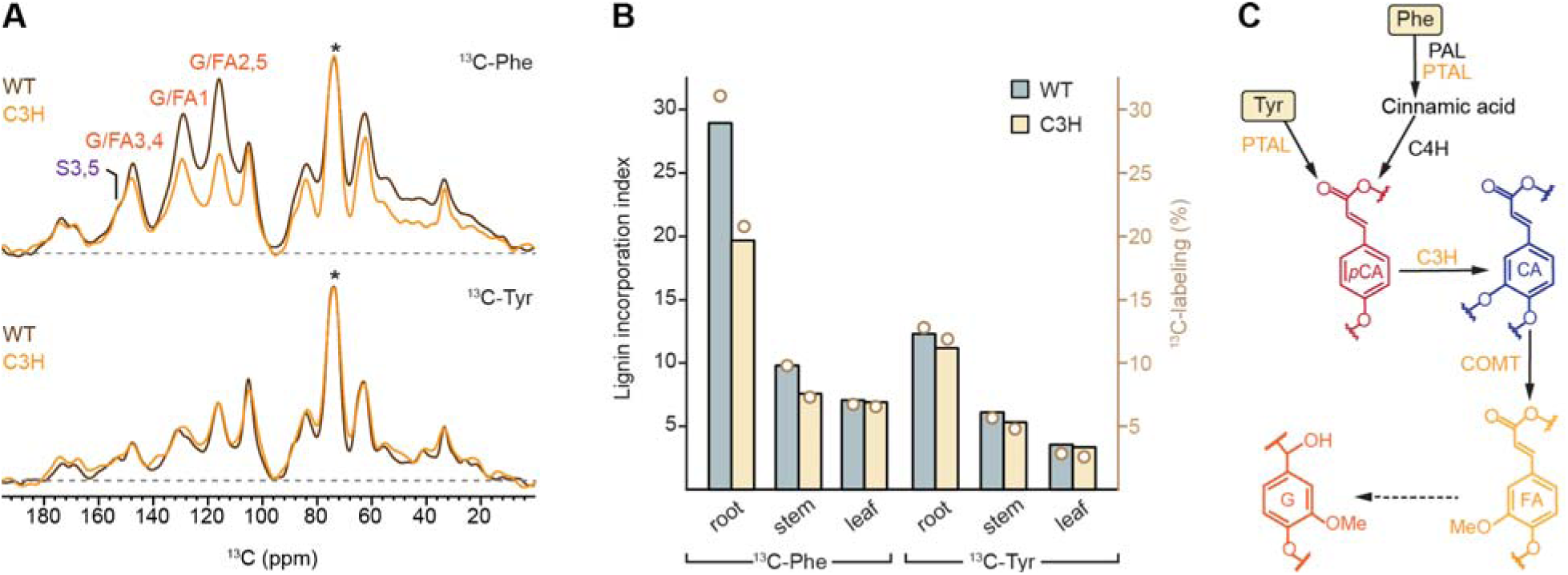
Impact of C3H knockdown on lignin biosynthesis from precursors in *Brachypodium* root. (**A**) Overlay of 1D ^13^C CP spectra from ^13^C-Phe-labeled (top) and ^13^C-Tyr-labeled (bottom) root samples for WT (black) and C3H-knockdown plants (orange). The ^13^C-Phe-labeled mutant shows a marked reduction in lignin-associated signal intensity. Asterisk: the highest peak of unlabeled carbohydrate used for spectral intensity normalization. (**B**) Bar graph comparing lignin incorporation indices in WT (frosted blue) and mutant (yellow) roots labeled with ^13^C-Phe (left) or ^13^C-Tyr (right). The open circles in camel represent the ^13^C-labeling percentage of each sample, projected to the y-axis on the right. (**C**) Simplified phenylpropanoid pathway highlighting the role of C3H in converting *p*-coumaric acid (*p*CA) to caffeic acid (CA), a key step channeling Phe-derived metabolites toward lignin biosynthesis. Phe enters upstream via PAL and proceeds through C3H, whereas Tyr enters via PTAL at *p*-coumarate, bypassing PAL, consistent with the reduced sensitivity of Tyr-derived lignin to the mutation.

In roots, the lignin incorporation index for phenylalanine decreased from 29 in wildtype plants to 20 in the C3H mutant, whereas the tyrosine-derived index remained largely unchanged at approximately 12 (**Figure 3B**). A similar, though less pronounced, preferential decline in ^13^C-Phe-labeled lignin was observed in stem tissues (**Figure S5**), where the index decreased from 10 to 8, while no consistent precursor-dependent differences were detected in leaves, consistent with their low lignin content (**Figure 3B**; **Figure S6**). These results indicate that the impact of C3H disruption on precursor incorporation is both precursor- and tissue-dependent.

This differential sensitivity can be explained by the distinct metabolic entry points of the two precursors within the phenylpropanoid pathway. Phenylalanine enters through the canonical route via PAL, proceeding through cinnamic acid and *p*-coumarate to caffeic acid before incorporation into lignin. In contrast, tyrosine enters downstream via PTAL directly at the level of *p*-coumarate (**Figure 3C**). This partial bypass of upstream steps reduces the dependence of tyrosine-derived lignin biosynthesis on C3H-mediated hydroxylation, explaining the relative resilience of tyrosine labeling to C3H disruption (50, 52). The C3H knockdown mutant was then analyzed using DNP, yielding sensitivity enhancements of 22-fold for the ^13^C-Phe-labeled root sample (**Figure 4A**) and 18-fold for the ^13^C-Tyr-labeled root sample (**Figure S3**). The resulting 2D ^13^C-^13^C correlation spectra revealed that disruption of C3H led to substantial remodeling of lignin composition rather than a uniform reduction in lignin content (**Figure 4B**).

**Figure 4.**
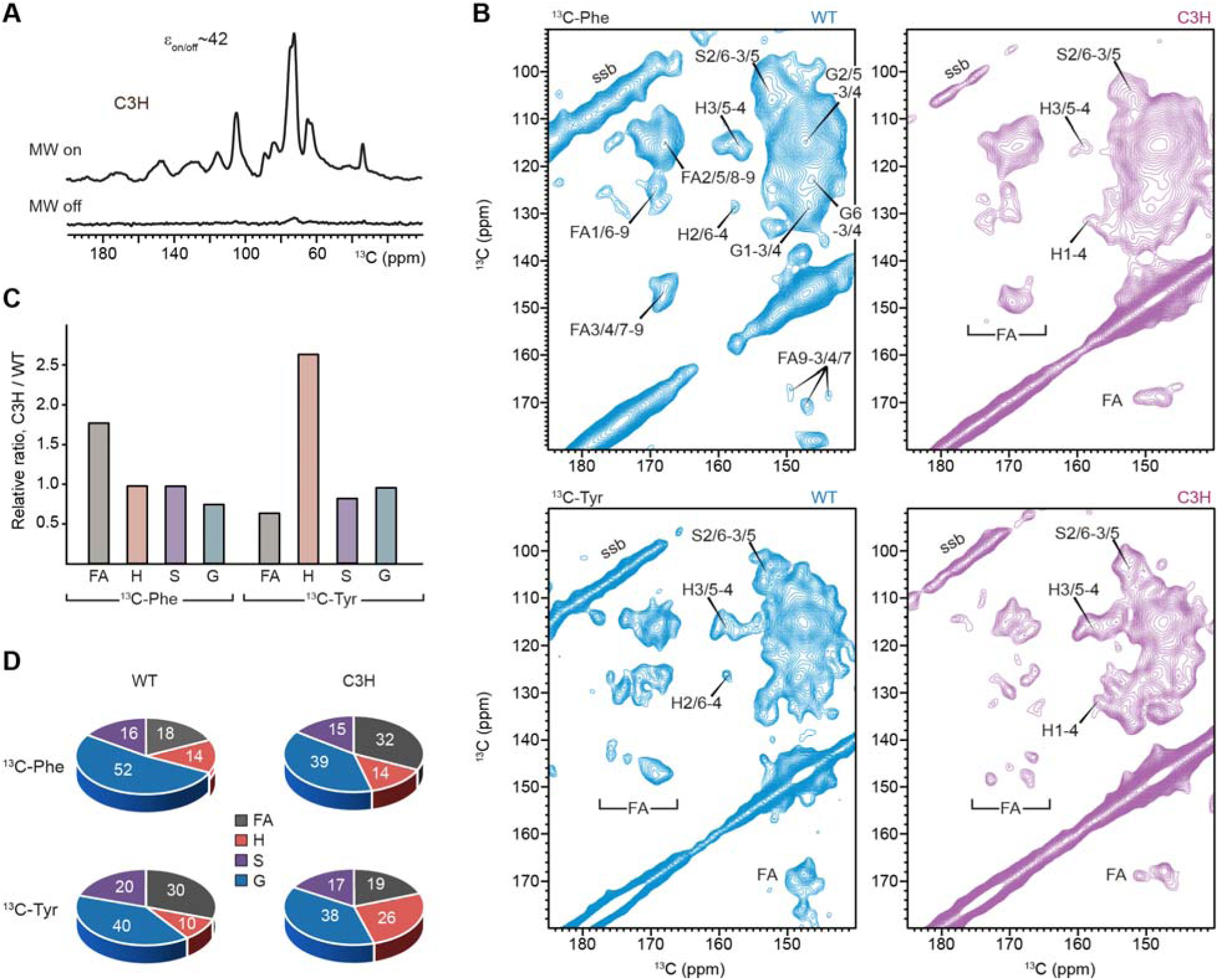
Altered lignin composition in C3H mutant revealed by DNP-enhanced 2D ^13^C-^13^C spectra. (**A**) DNP enhancement of NMR sensitivity in 1D ^13^C CP spectra of ^13^C-Phe-labeled C3H mutant roots acquired with (top) and without (bottom) microwave (MW) irradiation; the DNP enhancement factor (ε_on/off_) is 42. (**B**) DNP-enhanced 2D ^13^C-^13^C DARR spectra of WT (left, blue) and C3H (right, purple) roots labeled with ^13^C-Phe (top) or ^13^C-Tyr (bottom). Distinct aromatic and side-chain correlations from p-hydroxyphenyl (H), guaiacyl (G), syringyl (S), and ferulate (FA) units are observed. Spinning sideband: ssb. (**C**) Relative lignin-component ratios in the mutant for ^13^C-Phe-labeled (left) and ^13^C-Tyr-labeled (right) roots, expressed relative to WT. (**D**) Molar composition of lignin in ^13^C-Phe-labeled (top) and ^13^C-Tyr-labeled (bottom) roots of WT (left) and C3H (right); different lignin units are color-coded.

Notably, ferulate-derived signals responded in a precursor-dependent manner. In ^13^C-Phe-labeled roots, FA cross-peaks were substantially enhanced in the C3H mutant compared with wildtype, whereas in ^13^C-Tyr-labeled roots, FA signals were reduced upon C3H disruption (**Figure 4B**; **Figures S7**). Intensity analysis confirmed a 1.8-fold increase in FA content in the mutant when ^13^C-Phe was used as the precursor, but a decrease to 64% of wildtype levels when ^13^C-Tyr was used (**Figure 4C**). As a result, FA accounted for 18% of detected monolignol units in ^13^C-Phe-labeled wild-type roots but increased to 32% in the C3H mutant (**Figure 4D**). In contrast, FA content in ^13^C -Tyr-labeled roots decreased from 30% in wildtype to 19% in the mutant.

Spectral comparison further revealed a selective increase in H-lignin in the ^13^C-Tyr-labeled C3H mutant relative to the wild type, as evidenced by the enhanced H3/5-4 cross-peak (**Figure 4B**). Quantitative analysis showed that H-lignin intensity increased by approximately 2.6-fold in the mutant compared with wild type (**Figure 4C**), with its molar fraction rising from 10% to 26% (**Figure 4D**). These results indicate that, upon C3H disruption, tyrosine-derived intermediates are less efficiently retained within the hydroxycinnamate pool, specifically cell-wall-bound ferulates, and are instead redirected toward incorporation into H-type lignin.

In addition, G-lignin exhibited a moderate decline only in the ^13^C-Phe-labeled C3H mutant. In this condition, approximately three quarters of the G-lignin signal intensity was retained relative to the wildtype (**Figure 4C**), corresponding to a decrease in molar fraction from 52% to 39% (**Figure 4D**). No comparable reduction was observed in ^13^C-Tyr-labeled samples, in which the G-lignin fraction remained stable at ∼40% of total lignin. This precursor-specific trend mirrors the preferential loss of G-lignin signals observed for ^13^C-Phe labeling in the 1D ^13^C spectra (Figure 3A), suggesting that G-lignin formation is selectively sensitive to C3H disruption in the phenylalanine-derived pathway.

### Tyr- and Phe-driven lignification reveals pathway-specific control by C3H in grasses

Our results show that disruption of C3H elicits a precursor-dependent reorganization of lignin biosynthesis in *Brachypodium*. Phe-derived lignin is sensitive to C3H loss, showing reduced overall incorporation, overaccumulation of cell wall-bound FA and a concomitant decline in G-lignin. In contrast, Tyr-derived lignin exhibits a more buffered response to C3H disruption, maintaining overall incorporation into G-lignin and redistributing carbon away from free hydroxycinnamate pools toward H-lignin.

Overall, the solid-state NMR observations of lignin structure in C3H-deficient *Brachypodium* mutants grown under Tyr- and Phe-labeling conditions support a model in which lignification in grasses is modular and pathway-specific. These responses are best explained by a dual-route model of lignification in grasses, in which a Tyr-derived cytosolic soluble pathway and a Phe-derived canonical ER-localized pathway contribute differently to lignin and hydroxycinnamate biosynthesis (**Figure 5**) (50, 52). More specifically, phenylalanine is deaminated to cinnamic acid and converted to p-coumaroyl-CoA through the shikimate-ester pathway, which supports the formation of H-, G-, and S-lignin deposition via membrane-associated cytochrome P450 *p*-coumaroyl shikimate 3’-hydroxylase (C3′H) and F5H/COMT-dependent reactions. In contrast, tyrosine forms *p*-coumarate directly via PTAL and proceeds through a soluble route (or free-acid pathway) that requires the 3-hydroxylation of *p*CA to produce caffeic and ferulic acids.

**Figure 5.**
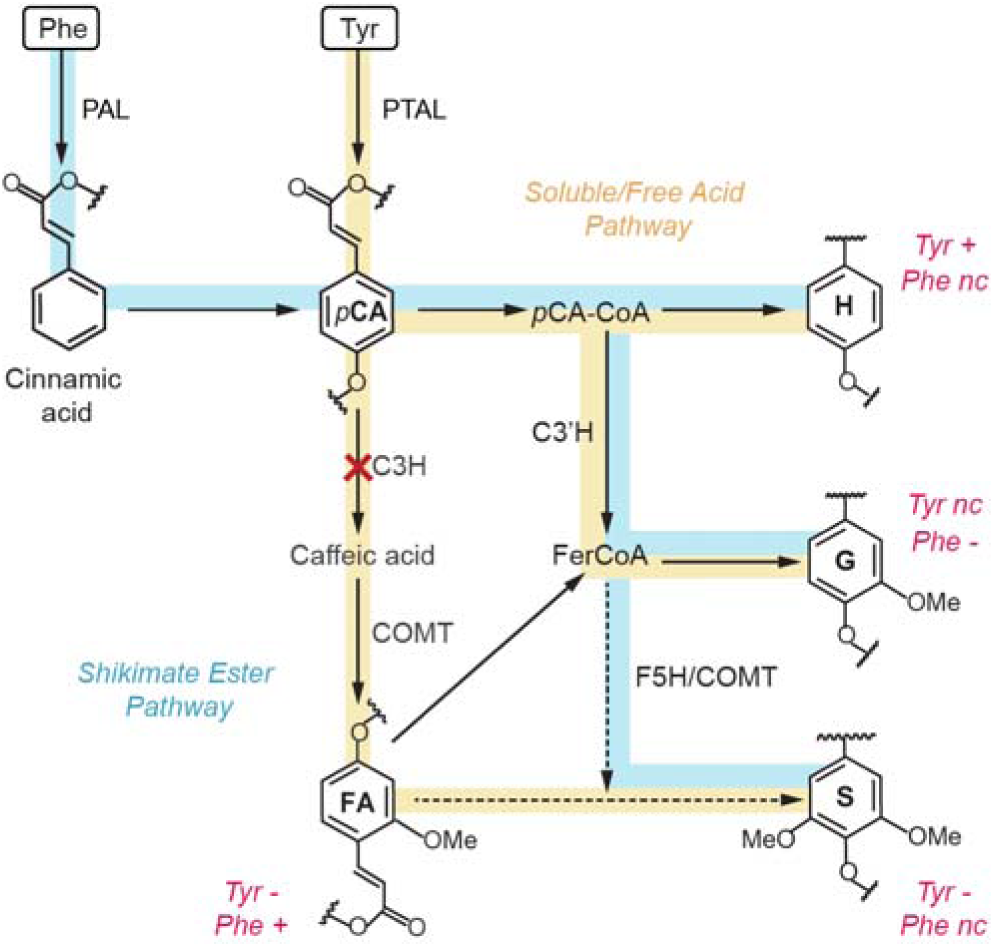
Distinct entry points of Phe and Tyr into phenylpropanoid metabolism. Schematic overview of the early phenylpropanoid steps focused on the two upstream routes. Phenylalanine enters the canonical ER-localized shikimate-ester pathway (blue), whereas tyrosine enters a soluble/free-acid pathway (beige). Solid-state NMR-observed content changes in Tyr- or Phe-labeled samples are annotated in magenta for an increase (+), decline (-), or no change (nc). Dash lines indicate paths where multiple reactions are needed.

Under C3H knockdown, the soluble pathway is impaired, and Tyr-derived *p*-CA cannot be efficiently 3-hydroxylated in the absence of functional C3H, leading to the increased incorporation of labeled Tyr into H-units (**Figure 5**). At the same time, the pools of ^13^C-Tyr required for FA and S-lignin formation are restricted. Notably, Tyr-derived G-lignin remained near wild-type levels, suggesting that C3H restricts flux mainly through the soluble pathway, with no impact on the shikimate ester route. In contrast, the PAL pathway remains active and Phe-derived phenylpropanoid flux continues to enter the canonical route. Reduced G-lignin deposition derived from the canonical pathway in the C3H lines can be explained by the ascorbate peroxidase activity of C3H, which may contribute to lignin polymerization in the cell wall (66). This may promote carbon repartitioning, consistent with the increased incorporation of ^13^C-Phe into the cell wall-bound FA pools. Moreover, the levels of ^13^C-Phe incorporated into S-lignin were consistent with buffering at downstream steps of the canonical pathway, likely via F5H and COMT branch, maintaining Phe-labeled S-lignin near wild-type levels despite perturbations in the free phenolic acids route. In addition to differences in subcellular localization, these parallel pathways may reflect cell type-specific contributions to lignification, with the soluble Tyr route providing hydroxycinnamates and S-lignin to fiber cells during stress-induced lignification, and the canonical Phe-derived route supporting developmental lignification in xylem vessels (12, 24).

## DISCUSSION

The results presented here establish that lignin biosynthesis in *Brachypodium distachyon* is strongly shaped by precursor identity, both under native conditions and in response to pathway perturbation. Precursor-resolved ^13^C-labeling combined with DNP-enhanced solid-state NMR demonstrates that phenylalanine and tyrosine contribute to lignin through distinct metabolic routes, conferring differential effects on polymer composition and robustness. These findings reveal that lignification in grasses is regulated not only by enzymatic capacity within the phenylpropanoid pathway, but also by the routing and utilization of specific aromatic amino acid pools.

Although metabolic flexibility in grass lignification has been recognized, the extent to which precursor identity itself governs lignin structure in planta has remained unclear (61, 67). By directly tracing precursor-derived carbon into intact cell walls, this study shows that phenylalanine is incorporated more efficiently into the lignin polymer (**Figure 1D, E**), whereas tyrosine contributes less to total lignin accumulation but preferentially enriches non-canonical subunits, including ferulate- and hydroxyphenyl-associated motifs (**Figure 2E**). This divergence establishes precursor entry point as a critical and previously underappreciated determinant of lignin heterogeneity in grasses (68).

These observed differences are consistent with the organization of the phenylpropanoid pathway in grasses. Phenylalanine enters through the canonical PAL-mediated route that supports high flux toward guaiacyl and syringyl monolignols, whereas tyrosine enters downstream via PTAL, a feature unique to grasses (18, 26, 69, 70). Although PTAL-mediated entry supports lignification, the lower overall incorporation of tyrosine-derived carbon suggests that this route functions as a secondary input that preferentially supplies structurally distinct lignin subunits rather than maximizing polymer yield.

A key implication of this aromatic amino acid precursor-level organization is revealed under pathway perturbation. Disruption of C3H does not simply reduce lignin deposition, but instead induces pronounced, precursor-dependent remodeling of lignin composition. Phenylalanine-derived lignin is highly sensitive to C3H loss, showing reduced guaiacyl incorporation and increased ferulate accumulation (**Figures 3A** and **4D**), whereas tyrosine-derived lignification remains comparatively resilient, maintaining overall polymer assembly despite marked shifts in subunit composition (**Figure 4C, D**).

This resilience highlights an important functional role for PTAL-mediated entry into the phenylpropanoid pathway. Because tyrosine enters at the level of *p*-coumarate, downstream of PAL and partially decoupled from upstream control points, tyrosine-derived flux is less constrained by C3H disruption (20, 52). Consequently, lignification can be sustained even when canonical monolignol biosynthesis is impaired, though with altered subunit composition, positioning tyrosine-mediated flux as a buffering mechanism that preserves cell-wall integrity under constrained phenylpropanoid flux (50, 71, 72).

The increased incorporation of hydroxyphenyl units in tyrosine-labeled C3H mutants further suggests that tyrosine-derived intermediates are redirected toward less substituted lignin structures under constrained flux. Such H-rich lignin has been associated with altered cross-linking and increased chemical accessibility, properties that may influence wall mechanics, stress responses, and biomass recalcitrance (73–75). From an evolutionary perspective, PTAL-mediated lignification may therefore represent an adaptive strategy that enhances cell-wall robustness under fluctuating metabolic or environmental conditions (4, 52).

Interpreting the precursor-dependent lignin remodeling observed here relies on recent advances in high-resolution solid-state NMR and an expanding body of lignocellulose data (76–78). Multidimensional ^13^C correlation experiments, fast magic-angle spinning ^1^H-detection, DNP enhancement, and selective polarization transfer techniques have revealed that cell-wall polymers are not randomly assembled but instead form a chemically and spatially organized network (79–85). In particular, xylan adopts a defined two-fold flat-ribbon conformation when bound to cellulose microfibrils and stabilizes lignin-polysaccharide interactions through its three-fold conformation, reframing the secondary cell wall as a regulated interpolymer network (36, 86–89).

This solid-state NMR and DNP approaches applied in the present study preserves native polymer context while enabling observation of precursor-derived carbon-carbon correlations in neo-synthesized and polymerized lignin. Labeling efficiencies reflect relative flux rather than absolute composition, and extension to additional tissues, developmental stages, or environmental conditions will further test the generality of precursor-dependent lignification. At the same time, we could not detect signals of tricin, which is typically reported to be incorporated as a lignin end group in grasses. This may be due to its low abundance in *Brachypodium*, accounting for only 0.5 wt% even in the aerial tissues where tricin is more abundant (23, 90), which makes it difficult to detect by solid-state NMR; therefore, follow-up studies are required to evaluate its role.

More broadly, the findings demonstrate that lignin composition in grasses is governed not only by enzyme activity but also by the availability and routing of precursor pools. This precursor-level control provides an additional lever for modulating lignin architecture and suggests new strategies for engineering biomass with improved processing characteristics (91, 92). By revealing tyrosine-mediated lignification as a compensatory and flexible pathway, this work advances our understanding of grass cell-wall metabolism and highlights metabolic plasticity as a key feature of plant structural biopolymers.

## MATERIALS AND METHODS

### Wildtype and mutant plants of *Brachypodium distachyon*

*Brachypodium distachyon* T-DNA line JJ25124 (IL000024891) was obtained from the Joint Genome Institute (JGI) T-DNA collection and corresponds to an activation-tagged line generated using the pJJ2LBA vector (50). In this line, the T-DNA is inserted in the last intron of the c3h gene (Bradi1g65820), in the positive orientation and located 121 bp upstream of the stop codon resulting in a partial loss of function (knockdown) mutant. The *Brachypodium distachyon* accession Bd213, the parental background of the T-DNA mutant population, was used as the wildtype control.

### Isotopic labeling and growth conditions

To trace carbon incorporation into lignin biosynthesis, *Brachypodium distachyon* seedlings were grown under isotopic labeling conditions using ^13^C-labeled precursors, following approaches previously developed for grasses (18, 50). Seeds were surface-sterilized and plated on half-strength Murashige and Skoog (MS) medium lacking sucrose, as it will likely dilute the incorporation of 13C-glucose, solidified with 0.5% Gelrite (pH 5.7), and grown for 5 weeks under continuous light conditions (120-150□μmol□m^-2^s^-1^) at 26□°C. The MS medium was supplemented with the following labeling treatments: i) 10 mM unlabeled D-glucose as a control (unlabeled), ii) 0.1 mM ^13^C -labeled phenylalanine plus 10 mM unlabeled glucose (Phe-labeled), iii) 0.1 mM ^13^C_9_-labeled tyrosine plus 10 mM unlabeled glucose (Tyr-labeled), iv) 10 mM ^13^C_6_-labeled glucose (Glc-labeled), or v) a combination of 0.1 mM ^13^C_9_-phenylalanine, 0.1 mM ^13^C_9_-tyrosine, and 10 mM ^13^C_6_-glucose (uniformly labeled).

All labeling stock media were prepared fresh. Specifically, ^13^C_9_-labeled Tyr and Phe (10 mM) were dissolved in a mixture of DMSO, HCl, and Milli-Q water at a ratio of 1:0.1:8.9 (v/v), and ^13^C_6_-glucose (1 M) was prepared in Milli-Q water. Approximately 15 seeds were plated per tube. After growth, roots, stems, and leaves were harvested, flash-frozen in liquid nitrogen, and stored at -80°C. For each sample, approximately 10 mg of material was packed into 3.2-mm Bruker MAS rotors for solid-state NMR measurements, and the remaining material was embedded in a DNP matrix and transferred to 3.2-mm sapphire rotors for MAS-DNP experiments.

### 1D ^13^C Solid-state NMR experiments

Solid-state NMR experiments were conducted using a Bruker Avance Neo spectrometer equipped with a 3.2-mm MAS HCN triple-resonance probe. The spectrometer operated at a magnetic field strength of 600 MHz (14.1 Tesla) at the Max T. Rogers Facility at Michigan State University. Experiments were performed at a MAS frequency of 14 kHz and a thermo-couple reported sample temperature of 283 K. ^13^C chemical shifts were externally referenced to tetramethylsilane (TMS) by calibrating the adamantane CH_2_ peak to 38.48 ppm (93), with the resulting spectral reference applied to the spectra collected on plant samples. Typical radiofrequency field strengths were 83 kHz for ^1^H decoupling and 83.3 kHz and 50-62.5 kHz for the 90° hard pulses of ^1^H and ^13^C, respectively. Due to the limited sample quantity and selective labeling, only 1D ^13^C NMR spectra could be acquired. 1D ^1^H-^13^C CP experiments were measured on all samples to selectively detect rigid molecular components. Each 1D ^13^C CP spectrum was acquired with 15,360 scans and a recycle delay of 2 s, resulting in a total acquisition time of approximately 8.5 h per sample. The key parameters and conditions of NMR experiments are listed in **Table S1**.

### MAS-DNP experiments

After acquisition of 1D ^13^C CP spectra using conventional solid-state NMR, the root samples labeled with ^13^C_9_-Phe and ^13^C_9_-Tyr were further processed for sensitivity-enhanced MAS-DNP measurements to enable 2D ^13^C-^13^C correlation experiments on these low-quantity samples. For each sample, approximately 10 mg of plant material was impregnated with 10 mM Asympol-POK (59, 94) in 50 μL of a cryoprotectant solvent mixture, referred to as the DNP matrix (or DNP juice), consisting of d_8_-glycerol/D_2_O/H_2_O at 60/30/10 Vol% (46, 58, 95). The samples were then manually ground in a chilled mortar and pestle for 15-20 min to promote radical diffusion into the plant cell wall material (96). The hydrated samples were subsequently packed into a 3.2-mm sapphire MAS rotor and sealed with silicone plugs for low-temperature MAS-DNP analysis.

DNP-enhanced solid-state NMR experiments were performed on a 600 MHz/395□GHz MAS-DNP spectrometer housed by National High Magnetic Field Laboratory (Tallahassee, FL) equipped with a 3.2□mm probe (97). Samples were spun at MAS frequencies of 10.5-11.2□kHz. The gyrotron microwave source operated at 395□GHz with a cathode current of 150□mA, a voltage of 16.2 kV, leading to approximatively 14.5 W of MW irradiation. The sample temperature was maintained at 110□K under microwave irradiation. The DNP signal buildup time constants ranaged from 0.6 s to 1.7 s. DNP enhancements were initially low for the mutant prepared using the original protocol but increased reproducibly after protocol optimization for the later experiments conducted. Because these measurements were obtained using different preparation workflows, the increase is attributed primarily to improved sample preparation. Using only the optimized protocol for comparisons, wild type and mutant showed similar enhancements within each sample-set condition. 2D ^13^C-^13^C correlation spectra correlation spectra were acquired using a 100 ms ^13^C-^13^C DARR mixing period, with 64 scans and a recycle delay of 1 s, resulting in a total acquisition time of approximately 16 h per sample. The ^13^C chemical shifts resolved for key lignin units were documented in **Table S2**.

### Estimation of precursor incorporation

To assess the extent of incorporation of ^13^C-labeled aromatic precursors into lignin, we analyzed 1D ^13^C CP spectra obtained from Phe- and Tyr-labeled root samples alongside an unlabeled control. The spectral region corresponding to aromatic lignin carbons was defined as 108-165 ppm, encompassing chemical shifts characteristic of monolignol subunits (13, 38). The total integral of this region was recorded for each spectrum. To account for variations in spectral scaling arising from differences in sample mass, acquisition parameters, or contact times, each spectrum was normalized to its highest-intensity peak, typically located at 72-73 ppm and corresponding to carbohydrate iC2/C3/C5 signals (57). We then defined a lignin incorporation index, calculated by comparing the integrated aromatic-region signal of a labeled sample to that of an unlabeled control and multiplying this ratio by a normalization factor that corrects for overall spectral intensity differences. Although not an absolute measure of lignin content, this index enables comparative assessment of precursor utilization efficiency across experimental conditions. These indices reflect relative incorporation trends rather than absolute carbon content; they represent how many-fold more lignin carbons are ^13^C-labeled from precursor incorporation compared with the unlabeled sample. The lignin-incorporation index was further converted to a ^13^C-labeling percentage by accounting for the contribution of naturally abundant ^13^C present in unlabeled molecules, with the procedures detailed in **Supplementary Text**.

## ACKNOWLDGMENT

The solid-state NMR analyses were supported by the U.S. Department of Energy under grant no. DE-SC0023702 to T.W. Sample labeling was supported by the University of Missouri start-up funds to J.B.A. A portion of this work was performed at the National High Magnetic Field Laboratory, which is supported by National Science Foundation Cooperative Agreement No. DMR-2128556 and the State of Florida. The MAS-DNP system at NHMFL is funded in part by NIH RM1-GM148766. F.J.S. was funded by the postdoctoral scholar award from the Provost’s Office at Florida State University.

## REFERENCES

1. C. Somerville, H. Youngs, C. Taylor, S. C. Davis, S. P. Long, Feedstocks for Lignocellulosic Biofuels. Science 329, 790–792 (2010).

2. D. B. Sulis et al., Advances in lignocellulosic feedstocks for bioenergy and bioproducts. Nat. Commun. 16, 1244 (2025).

3. W. Boerjan, J. Ralph, M. Baucher, Lignin biosynthesis. Annu. Rev. Plant Biol. 54, 519–546 (2003).

4. J. K. Weng, C. Chapple, The origin and evolution of lignin biosynthesis. New Phytol. 187, 273–285 (2010).

5. E. Novo-Uzal, F. Pomar, L. V. Gomez Ros, J. M. Espineira, A. R. Barcelo, Evolutionary History of Lignins. Adv. Bot. Res. 61, 311–350 (2012).

6. A. Zoghlami, G. Paes, Lignocellulosic Biomass: Understanding Recalcitrance and Predicting Hydrolysis. Front. Chem. 7, 874 (2019).

7. A. J. Rgauskas et al., Lignin Valorization: Improving Lignin Processing in the Biorefinery. Science 344, 6185 (2014).

8. M. E. Himmel et al., Biomass recalcitrance: engineering plants and enzymes for biofuels production. Science 315, 804–807 (2007).

9. R. A. Dixon, J. Barros, Lignin biosynthesis: old roads revisited and new roads explored. Open Biol. 9, 190215 (2019).

10. R. Vanholme, B. Demedts, K. Morreel, J. Ralph, W. Boerjan, Lignin biosynthesis and structure. Plant Physiol 153, 895–905 (2010).

11. L. M. Peracchi, R. Panahabadi, J. Barros-Rios, L. E. Bartley, K. A. Sanguinet, Grass lignin: biosynthesis, biological roles, and industrial applications. Front. Plant Sci. 15, 1343097 (2024).

12. J. Barros, H. Serk, I. Granlund, E. Pesquet, The cell biology of lignification in higher plants. Ann. Bot. 115, 1053–1074 (2015).

13. J. Ralph et al., Lignins: natural polymers from oxidative coupling of 4-hydroxyphenyl-propanoids. Phytochem. Rev. 3, 29–60 (2004).

14. Y. Tobimatsu, M. Schuetz, Lignin polymerization: how do plants manage the chemistry so well? Curr. Opin. Biotechnol. 56, 75–81 (2019).

15. R. Sibout et al., Structural redesigning Arabidopsis lignins into alkali-soluble lignins through the expression of p-coumaroyl-CoA: monolignol transferase PMT. Plant Physiol 170, 1358–1366 (2016).

16. R. Sibout et al., CINNAMYL ALCOHOL DEHYDROGENASE-C and-D are the primary genes involved in lignin biosynthesis in the floral stem of Arabidopsis. Plant Cell 17, 2059–2076 (2005).

17. P. Wang et al., A 13C isotope labeling method for the measurement of lignin metabolic flux in Arabidopsis stems. Plant Methods 14, 51 (2018).

18. J. Barros et al., Role of bifunctional ammonia-lyase in grass cell wall biosynthesis. Nat. Plants 2, 1–9 (2016).

19. J. Rosler, F. Krekel, N. Amrhein, J. Schmid, Maize phenylalanine ammonia-lyase has tyrosine ammonia-lyase activity. Plant Physiol. 113, 175–179 (1997).

20. H. A. Maeda, Lignin biosynthesis: Tyrosine shortcut in grasses. Nat. Plants 2, 1–2 (2016).

21. V. B. Caroline et al., Metabolic engineering of a tyrosine-specific phenylpropanoid pathway in plants. BioRxiv, DOI: 10.64898/2025.12.16.694581 (2025).

22. W. Lan et al., Tricin, a Flavonoid Monomer in Monocot Lignification Plant Physiol. 167, 1284–1295 (2015).

23. W. Lan et al., Tricin-lignins: occurrence and quantitation of tricin in relation to phylogeny. Plant J. 88, 1046–1057 (2016).

24. W. Zhu, J. Barros, Tissue-Specific Developmental Changes in Lignin Deposition in Model Plants. *Physiol*. Planta. 177, e70607 (2025).

25. F. Chen, R. A. Dixon, Lignin modification improves fermentable sugar yields for biofuel production. Nat. Biotechnol 25, 759–761 (2007).

26. H. Maeda, N. Dudareva, The shikimate pathway and aromatic amino acid biosynthesis in plants. Annu. Rev. Plant Biol. 63, 73–105 (2012).

27. S. T. Withers, J. D. Keasling, Biosynthesis and engineering of isoprenoid small molecules. Appl. Microbiol. Biotechnol. 73, 980–990 (2007).

28. F. Ma, L. J. Jazmin, J. D. Young, D. K. Allen, Isotopically nonstationary 13C flux analysis of changes in Arabidopsis thaliana leaf metabolism due to high light acclimation. Proc. Natl. Acad. Sci. USA 111, 16967–16972 (2014).

29. X. Rao, J. Barros, Modeling lignin biosynthesis: a pathway to renewable chemicals. Trends Plant Sci. 29, 546–559 (2024).

30. J. Ralph, C. Lapierre, W. Boerjan, Lignin structure and its engineering. Curr. Opin. Biotechnol. 56, 240–249 (2019).

31. J. Barros et al., Proteomic and metabolic disturbances in lignin-modified Brachypodium distachyon. Plant Cell 34, 3339–3363 (2022).

32. B. Reif, S. E. Ashbrook, L. Emsley, M. Hong, Solid-State NMR Spectroscopy. Nat. Rev. Methods Primers 1, 2 (2021).

33. C. R. Munson, Y. Gao, J. C. Mortimer, M. D. T., Solid-State Nuclear Magnetic Resonance as a Tool to Probe the Impact of Mechanical Preprocessing on the Structure and Arrangement of Plant Cell Wall Polymers. Front. Plant. Sci. 12, 766506 (2022).

34. L. D. Fernando, W. Zhao, I. Gautam, A. Ankur, T. Wang, Polysacchride Assemblies in Fungal and Plant Cell Walls Explored by Solid-State NMR. Structure 31, 1375–1385 (2023).

35. P. Duan et al., Xylan Structure and Dynamics in Native Brachypodium Grass Cell Walls Investigated by Solid-State NMR Spectroscopy. ACS Omega 6, 15460–15471 (2021).

36. T. J. Simmons et al., Folding of xylan onto cellulose fibrils in plant cell walls revealed by solid-state NMR. Nat. Commun. 7, 13902 (2016).

37. Y. Gao, A. S. Lipton, Y. Wittmer, D. T. Murray, J. C. Mortimer, A grass-specific cellulose-xylan interaction dominates in sorghum secondary cell walls. Nat. Commun. 11, 6081 (2020).

38. X. Kang et al., Lignin-polysaccharide interactions in plant secondary cell walls revealed by solid-state NMR. Nat. Commun. 10, 347 (2019).

39. Q. Z. Ni et al., High Frequency Dynamic Nuclear Polarization. Acc. Chem. Res. 46, 1933–1941 (2013).

40. A. J. Rossini et al., Dynamic Nuclear Polarization Surface Enhanced NMR Spectroscopy. Acc. Chem. Res. 46, 1942–1951 (2013).

41. K. R. Thurber, W. M. Yau, R. Tycko, Low-temperature dynamic nuclear polarization at 9.4 T with a 30 mW microwave source *J*. Magn. Reson. 204, 303–313 (2010).

42. F. Mentink-Vigier et al., Nuclear depolarization and absolute sensitivity in magic-angle spinning cross effect dynamic nuclear polarization. Phys. Chem. Chem. Phys. 17, 21824–21836 (2015).

43. W. Y. Chow, G. De Papae, S. Hediger, Biomolecular and biological applications of solid-state NMR with dynamic nuclear polarization enhancement. Chem. Rev. 122, 9795–9847 (2022).

44. T. Biedenba□nder, V. Aladin, S. Saeidpour, B. Corzilius, Dynamic nuclear polarization for sensitivity enhancement in biomolecular solid-state NMR. Chem. Rev. 122, 9738–9794 (2022).

45. W. Zhao et al., Enriched Molecular-Level View of Saline Wetland Soil Carbon by Sensitivity-Enhanced Solid-State NMR. J. Am. Chem. Soc. 147, 519–531 (2024).

46. H. Takahashi, D. Lee, L. Dubois, M. Bardet, S. Hediger, Rapid natural-abundance 2D 13C-13C correlation spectroscopy using dynamic nuclear polarization enhanced solid-state NMR and matrix-free sample preparation. Angew. Chem. Int. Ed. 124, 11936–11939 (2012).

47. F. A. Perras et al., Atomic-level structure characterization of biomass pre- and post-lignin treatment by dynamic nuclear polarization-enhanced solid-state NMR. J. Phys. Chem. A 121, 623–630 (2016).

48. J. Viger-Gravel et al., Topology of Pretreated Wood Fibers Using Dynamic Nuclear Polarization. J. Phys. Chem. C 123, 30407–30415 (2019).

49. J. Draper et al., Brachypodium distachyon. A new model system for functional genomics in grasses. Plant Physiol. 127, 1539–1555 (2001).

50. J. Barros et al., 4-Coumarate 3-hydroxylase in the lignin biosynthesis pathway is a cytosolic ascorbate peroxidase. Nat. Commun 10, 1994 (2019).

51. J. E. Thompson, S. C. Fry, Density-labelling of cell wall polysaccharides in cultured rose cells: comparison of incorporation of 2H and 13C from exogenous glucose. Carbohydr. Res. 332, 175–182 (2001).

52. J. Barros, R. A. Dixon, Plant phenylalanine/tyrosine ammonia-lyases. Trends Plant Sci. 25, 66–79 (2020).

53. N. C. Carpita, Structure and biogenesis of the cell walls of grasses. Annu. Rev. Plant Biol. 47, 445–476 (1996).

54. H. V. Scheller, P. Ulvskov, Hemicelluloses. Annu. Rev. Plant Biol. 61, 263–289 (2010).

55. J. Vogel, Unique aspects of the grass cell wall. Curr. Opin. Biotechnol. 11, 301–307 (2008).

56. S. E. Sattler, D. L. Funnell-Harris, Modifying lignin to improve bioenergy feedstocks: strengthening the barrier against pathogens? Front. Plant Sci. 4, 70 (2013).

57. W. Zhao et al., Solid-state NMR of unlabeled plant cell walls: high-resolution structural analysis without isotopic enrichment. Biotechnol. Biofuels 14, 14 (2021).

58. C. Sauvée et al., Highly efficient, water□soluble polarizing agents for dynamic nuclear polarization at high frequency. Angew. Chem. Int. Ed. 52, 10858–10861 (2013).

59. F. Mentink-Vigier et al., Computationally Assisted Design of Polarizing Agents for Dynamic Nuclear Polarization Enhanced NMR: The AsymPol Family. J. Am. Chem. Soc. 140, 11013–11019 (2018).

60. J. Ralph, R. D. Hatfield, Pyrolysis-GC-MS characterization of forage materials. J. Agric. Food Chem. 39, 1426–1437 (1991).

61. R. D. Hatfield et al., Grass lignin acylation: p-coumaroyl transferase activity and cell wall characteristics of C3 and C4 grasses. Planta 229, 1253–1267 (2009).

62. R. D. Hatfield, J. Ralph, J. H. Grabber, Cell wall cross□linking by ferulates and diferulates in grasses. J. Sci. Food Agric. 79, 403–407 (1999).

63. H. B. Molinari, T. K. Pellny, J. Freeman, P. R. Shewry, R. A. Mitchell, Grass cell wall feruloylation: distribution of bound ferulate and candidate gene expression in Brachypodium distachyon. Front. Plant Sci. 4, 50 (2013).

64. W. Xiong et al., Mutation of 4-coumarate: coenzyme A ligase 1 gene affects lignin biosynthesis and increases the cell wall digestibility in maize brown midrib5 mutants. Biotechnol. Biofuels 12, 82 (2019).

65. Y. Deng, S. Lu, Biosynthesis and regulation of phenylpropanoids in plants. Crit. Rev. Plant Sci. 36, 257–290 (2017).

66. J. Zhang et al., PtomtAPX is an autonomous lignification peroxidase during the earliest stage of secondary wall formation in Populus tomentosa Carr. Nat. Plants 8, 828–839 (2022).

67. R. Vanholme et al., Metabolic engineering of novel lignin in biomass crops New Phytol. 196, 978–1000 (2012).

68. J. El-Azaz et al., Coordinated regulation of the entry and exit steps of aromatic amino acid biosynthesis supports the dual lignin pathway in grasses. Nat. Commun. 14, 7242 (2023).

69. T. Vogt, Phenylpropanoid biosynthesis. Mol. Plant 3, 2–20 (2010).

70. S. Supatmi et al., Essential yet dispensable: The role of CINNAMATE 4-HYDROXYLASE in rice cell wall lignification Plant Physiol. 198, kiaf164 (2025).

71. J. Ralph, T. Akiyama, H. D. Coleman, S. D. Mansfield, Effects on Lignin Structure of Coumarate 3-Hydroxylase Downregulation in Poplar. Bioenergy Res. 5, 1009–1019 (2012).

72. J. P. Simpson, J. Olson, B. Dilkes, C. Chapple, Identification of the Tyrosine- and Phenylalanine-Derived Soluble Metabolomes of Sorghum. Front. Plant Sci. 12, 714164 (2021).

73. M. Li, Y. Pu, A. J. Ragauskas, Current Understanding of the Correlation of Lignin Structure with Biomass Recalcitrance. Front. Chem. 4, 45 (2016).

74. M. Ozparpucu et al., Unravelling the impact of lignin on cell wall mechanics: a comprehensive study on young poplar trees downregulated for CINNAMYL ALCOHOL DEHYDROGENASE (CAD) Plant J. 91, 480–490 (2017).

75. E. Pesquet, I. Cesarino, S. Kajita, K. Pawlowski, Physiological roles of lignins – tuning cell wall hygroscopy and biomechanics. New Phytol. 248, 2674–2706 (2025).

76. N. Ghassemi et al., Solid-State NMR Investigations of Extracellular Matrixes and Cell Walls of Algae, Bacteria, Fungi, and Plants. Chem. Rev. 122, 10036–10086 (2022).

77. P. Xiao et al., Revealing structure and shaping priorities in plant and fungal cell wall architecture via solid-state NMR. Cell Surf. 14, 100159 (2025).

78. D. Debnath et al., Structure-guided utilization of lignocellulose for catalysis, energy, and biomaterials. Cell Rep. Phys. Sci. 6, 102911 (2025).

79. P. Xiao et al., Rapid High-Resolution Analysis of Polysaccharide-Lignin Interactions in Secondary Plant Cell Walls Using Proton-Detected Solid-State NMR. Anal. Chem. 97, 18046–18054 (2025).

80. B. Addison et al., Atomistic, macromolecular model of the Populus secondary cell wall informed by solid-state NMR. Sci. Adv. 10, adi7965 (2024).

81. T. Le Marchand et al., 1H-Detected Biomolecular NMR under Fast Magic-Angle Spinning. Chem. Rev. 122, 9943–10018 (2022).

82. P. Phyo, M. Hong, Fast MAS 1H–13C correlation NMR for structural investigations of plant cell walls. J. Biomol. NMR 73, 661–674 (2019).

83. P. Duan, M. Hong, Selective Detection of Intermediate-Amplitude Motion by Solid-State NMR. J. Phys. Chem. B 128, 2293–2303 (2024).

84. F. Deligey et al., Structure of In Vitro-Synthesized Cellulose Fibrils Viewed by Cryo-Electron Tomography and 13C Natural-Abundance Dynamic Nuclear Polarization Solid-State NMR. Biomacromolecules 23, 2290–2301 (2022).

85. B. Addison, M. C. Dickwella Widanage, Y. Pu, A. J. Ragauskas, A. E. Harman-Ware, Solid-state NMR at natural isotopic abundance for bioenergy applications. Biotechnol. Biofuels Bioprod. 46, 46 (2025).

86. O. M. Terrett et al., Molecular architecture of softwood revealed by solid-state NMR. Nat. Commun. 10, 4978 (2019).

87. P. Xiao et al., Emergence of lignin-carbohydrate interactions during plant stem maturation visualized by solid-state NMR. Nat. Commun. 16, 8010 (2025).

88. A. Kirui et al., Carbohydrate-aromatic interface and molecular architecture of lignocellulose. Nat. Commun. 13, 538 (2022).

89. Y. Yoshimi et al., Glucomannan engineering highlights roles of galactosyl modification in fine-tuning cellulose-glucomannan interaction in Arabidopsis cell walls. Nat. Commun. 16, 1235 (2025).

90. F. Chen, C. Zhuo, X. Xiao, T. H. Pendergast, K. M. Devos, A rapid thioacidolysis method for biomass lignin composition and tricin analysis. Biotechnol. Biofuels 14, 18 (2021).

91. N. D. Bonawitz, C. Chapple, The genetics of lignin biosynthesis: connecting genotype to phenotype. Annu. Rev. Genet. 44, 337–363 (2010).

92. N. D. Bonawitz, C. Chapple, Can genetic engineering of lignin deposition be accomplished without an unacceptable yield penalty? Curr. Opin. Biotechnol. 24, 336–343 (2013).

93. C. R. Morcombe, K. W. Zilm, Chemical shift referencing in MAS solid state NMR. J. Magn. Reson. 162, 479–486 (2003).

94. R. Harrabi et al., Highly Efficient Polarizing Agents for MAS-DNP of Proton-Dense Molecular Solids. Angew. Chem. Int. Ed. 61, e202114103 (2022).

95. A. Kumar et al., Optimizing chemistry at the surface of prodrug-loaded cellulose nanofibrils with MAS-DNP. Commun. Chem. 6, 58 (2023).

96. A. Kirui et al., Atomic resolution of cotton cellulose structure enabled by dynamic nuclear polarization solid-state NMR. Cellulose 26, 329–339 (2019).

97. M. Rosay et al., Solid-state dynamic nuclear polarization at 263 GHz: spectrometer design and experimental results. Phys. Chem. Chem. Phys. 12, 5850–5860 (2010).

