## Supplementary files for "Precursor-Dependent Routing of Aromatic Amino Acids Determines Lignin Structure in Grasses by Sensitivity-Enhanced Solid-State NMR"

This PDF file includes:

Supporting text

Figures S1 to S7

Tables S1 to S2

SI References

**Supporting Text**

**Estimation of ^13^C Lignin Enrichment.** Lignin ^13^C enrichment following [^13^C_9_-phenylalanine /^13^C_9_-tyrosine] feeding was quantified using lignin incorporation index (I), defined as the fold increase in aromatic-region ^13^C solid-state NMR signal intensity in labeled samples relative to unlabeled controls. Aromatic ^13^C signal intensities were normalized prior to comparison. The measured incorporation index reflects contributions from both isotopically introduced ^13^C and naturally occurring ^13^C, which is present at natural abundance in all samples. To determine the fraction of lignin carbon originating specifically from labeled phenylalanine or tyrosine, the contribution from natural abundance ^13^C was explicitly accounted for.

For labeled samples, the lignin ^13^C fraction is given by

$$f_{\text{lab}}=x+(1-x)\cdot0.011$$

where *x* denotes the fraction of lignin carbon atoms derived from the uniformly ^13^C labeled precursor and 0.011 corresponds to the natural abundance ^13^C fraction (1.1%). Unlabeled controls contain only natural abundance ^13^C (*f*_unl_ = 0.011). Accordingly, the lignin incorporation index is given by

$$I=\frac{f_{\text{lab}}}{f_{\text{unl}}}$$

From this relationship, the fraction of lignin carbon derived from labeled phenylalanine or tyrosine was calculated as

$$x=\frac{0.011(I-1)}{0.989}$$

The corresponding labeled carbon percentage can be obtained as

$$x(\%)=\frac{1.1(I-1)}{0.989}$$

This correction removes the contribution from natural abundance ^13^C and enables quantitative estimation of the fraction of lignin carbon derived from isotopically labeled aromatic amino acid precursors.

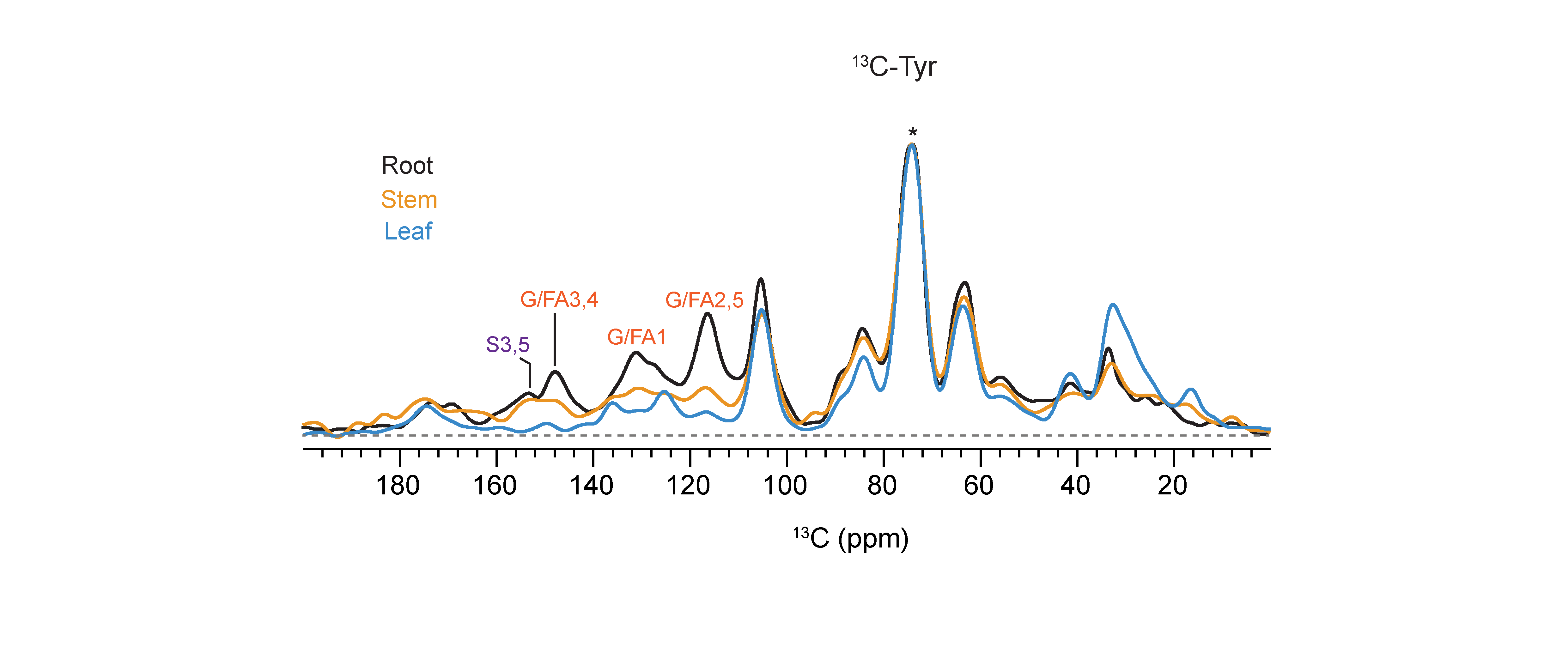

**Figure S1.** **Tissue-specific labeling efficiency across *Brachypodium* tissues.** (**A**) 1D ^13^C CP spectra of ^13^C-Tyr labeled root (black), stem (orange), and leaf (blue). The asterisk denotes the dominant carbohydrate peak at 73 ppm that is not ^13^C-enriched (from natural abundance of ^13^C present in unlabeled carbohydrates); this peak was used as the reference for intensity normalization. Lignin signals decrease sequentially from root to stem to leaf.

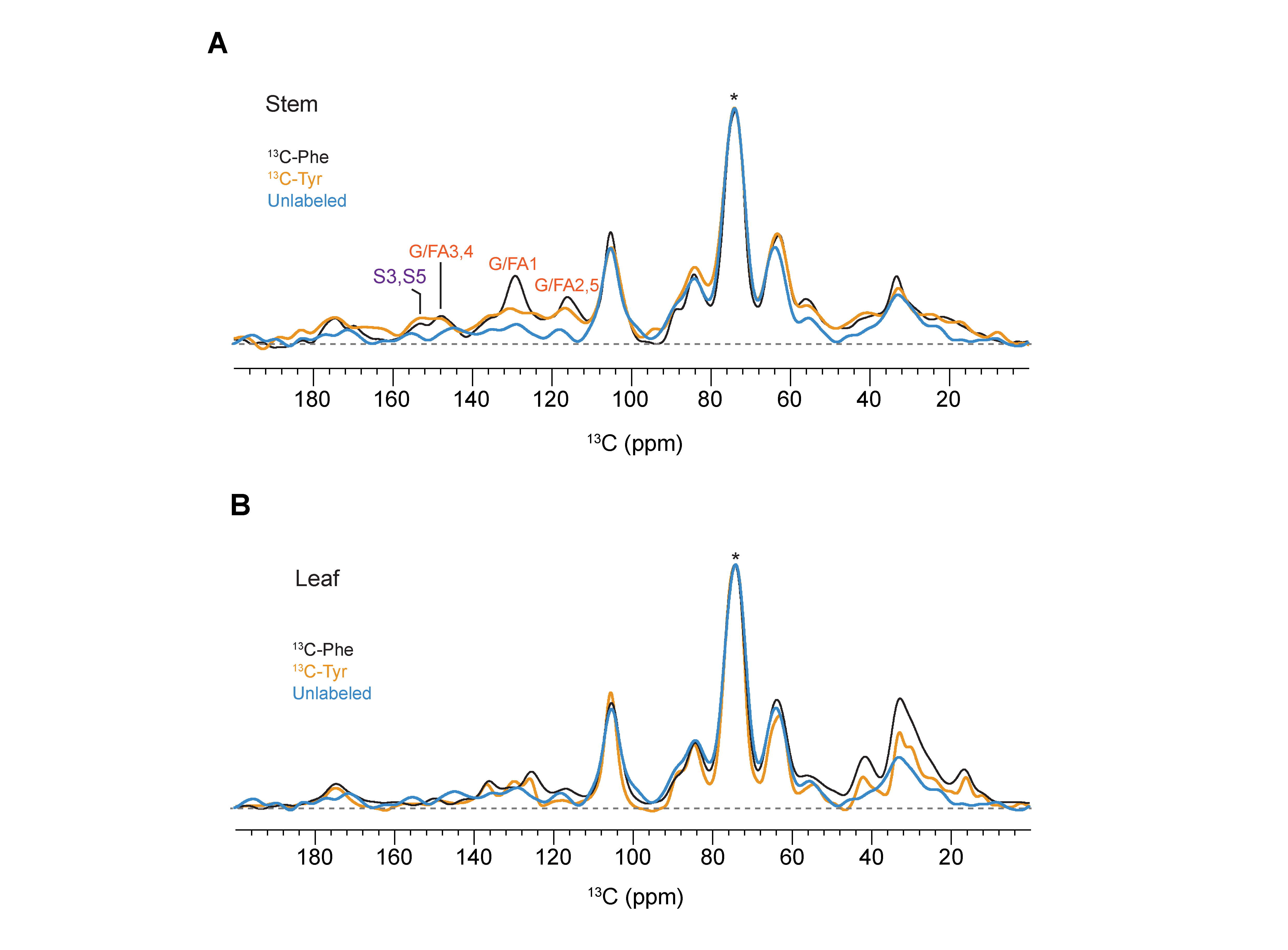

**Figure S2.** **Tissue-specific differences in lignin incorporation from precursors.** (**A**) Overlay of 1D ^13^C CP spectra collected on stem tissues enriched using ^13^C-Phe (blue), ^13^C-Tyr (Purple), alongside an Unlabeled control (blue). Labeling with ^13^C-Phe yields higher aromatic region signal intensity than ^13^C-Tyr, although the overall intensity is lower than that observed in root tissues. (**B**) Corresponding overlay for leaf tissues shows minimal lignin signal for both precursors, indicating weak lignification and negligible incorporation of labeled precursors.

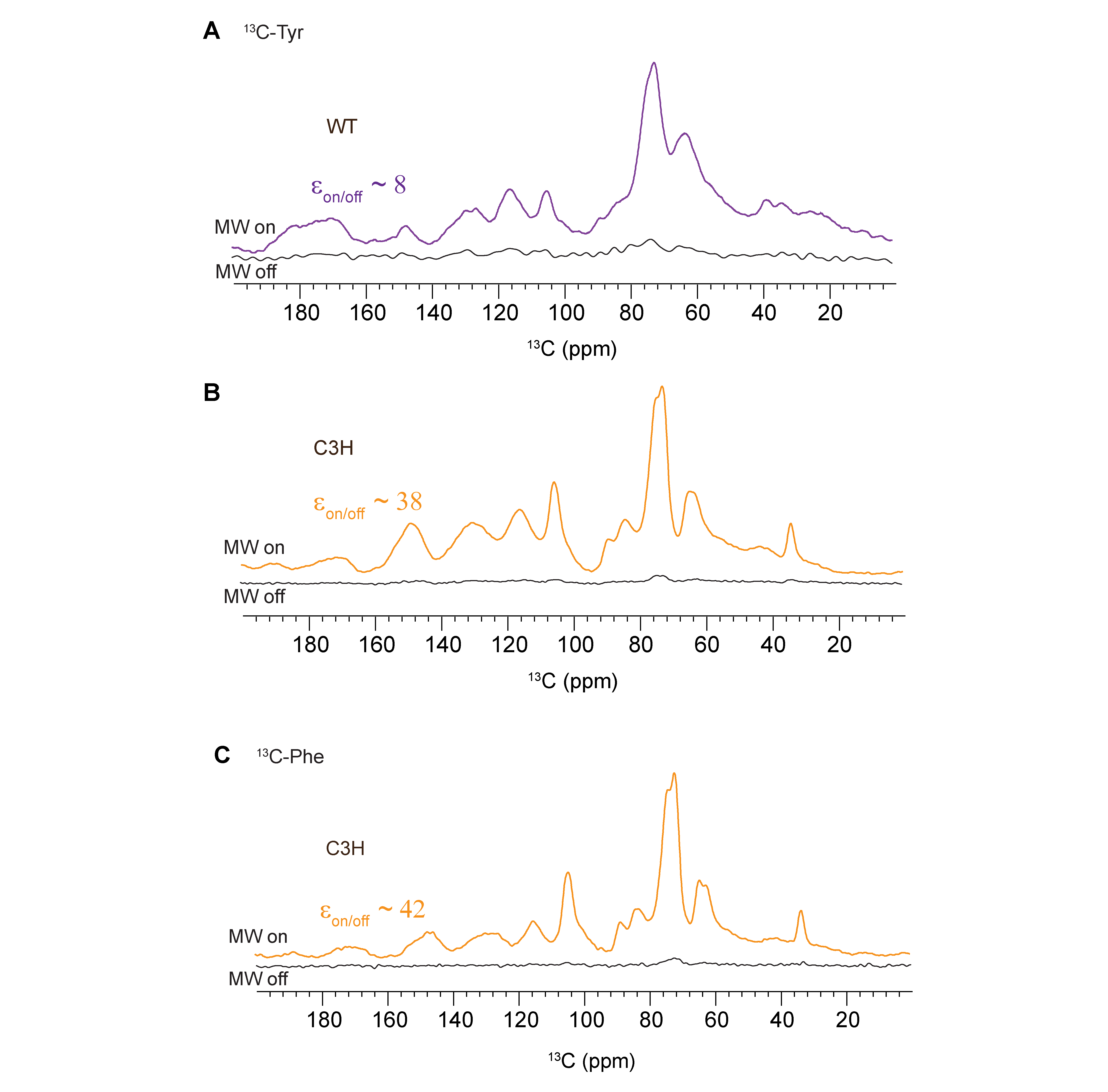

**Figure S3 Sensitivity enhanced DNP spectra of isotopically labeled root tissues.** (**A)** 1D ^13^C CP spectra of ^13^C-Tyr-labeled WT roots acquired with microwave irradiation (top) and without microwave irradiation (bottom), showing an enhancement factor (ε_on/off_) of 8. (**B**) Corresponding 1D ^13^C CP spectra of ^13^C-Tyr-labeled C3H roots acquired with microwave irradiation (top) and without microwave irradiation (bottom), showing an enhancement factor (ε_on/off_) of 38. (**C**) 1D ^13^C CP spectra of ^13^C-Phe-labeled C3H roots acquired with microwave irradiation (top) and without microwave irradiation (bottom), showing an enhancement factor (ε_on/off_) of 42.

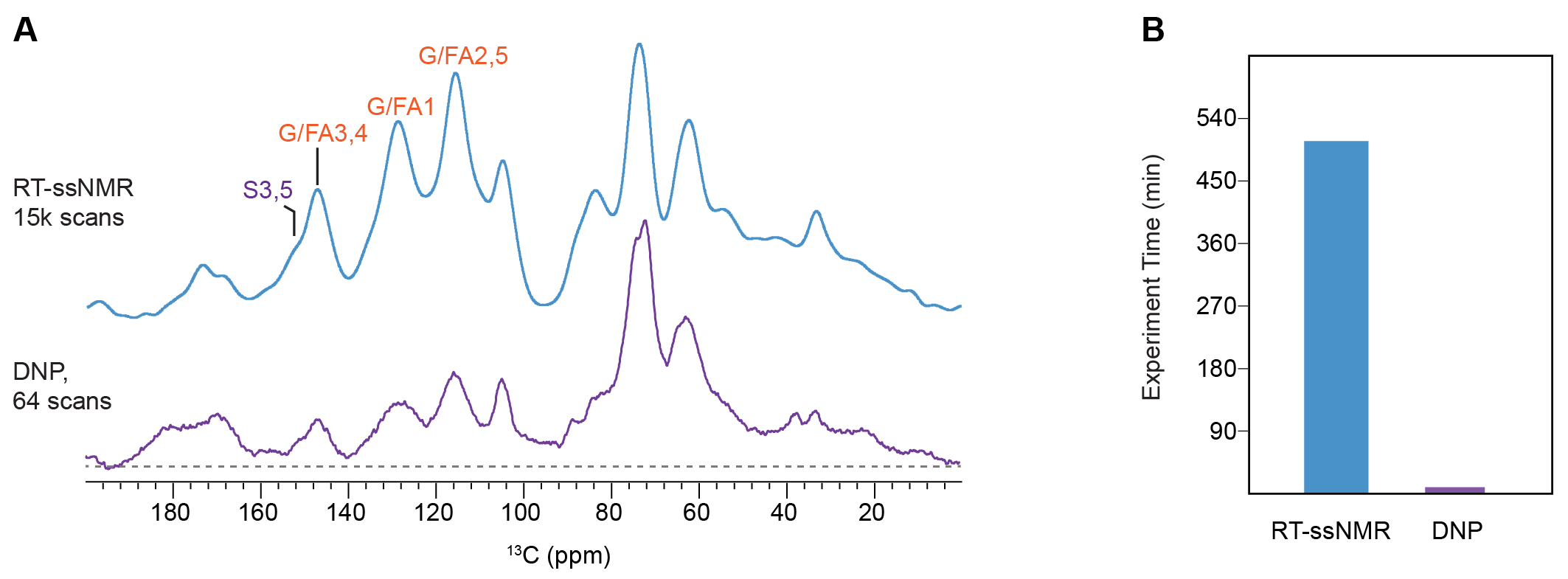

**Figure S4.** **Comparison of conventional CP and DNP enhanced CP spectra.** (**A**) Overlay of 1D ^13^C CP spectra acquired using conventional CP with 15,000 scans (blue) and DNP-enhanced CP with 64 scans (purple) for ^13^C-Phe labeled root tissue. The two spectra show nearly identical aromatic profiles, demonstrating that DNP enables comparable spectral quality with a dramatically reduced experiment time. (**B**) Histogram comparing the total experiment times, showing a reduction from 510 min for conventional CP to 1.5 min for DNP-enhanced CP.

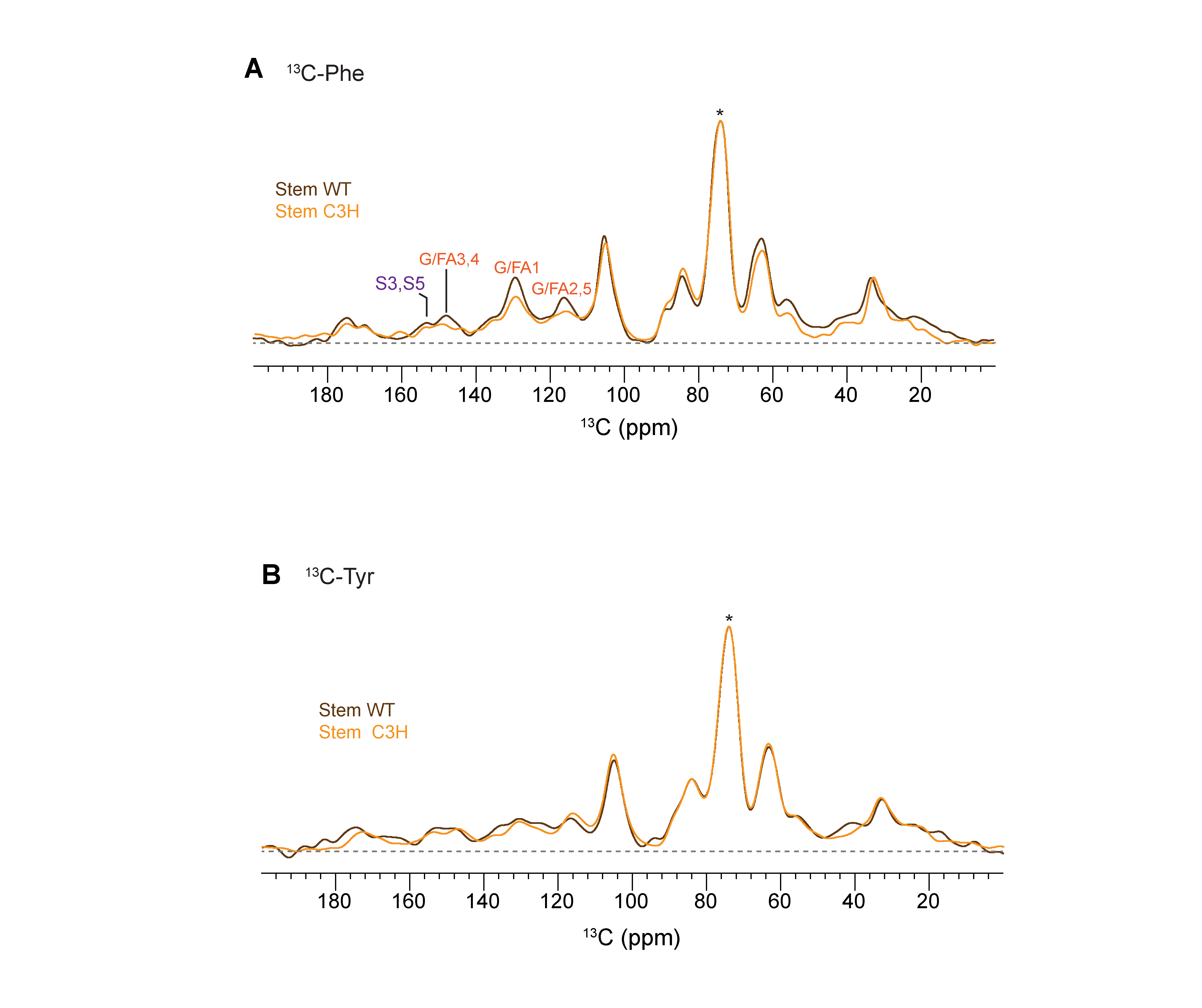

**Figure S5.** **Stem tissue response to C3H mutation under precursor labeling.** (**A**) CP-MAS spectra from ^13^C labeled WT (brown) and C3H (orange) mutant stem tissues show moderate reduction in aromatic signal upon mutation. (**B**)Corresponding spectra ^13^C-Tyr labeled stem tissues show negligible change. Compared to root, stem tissues show lower overall lignin incorporation and smaller mutant response.

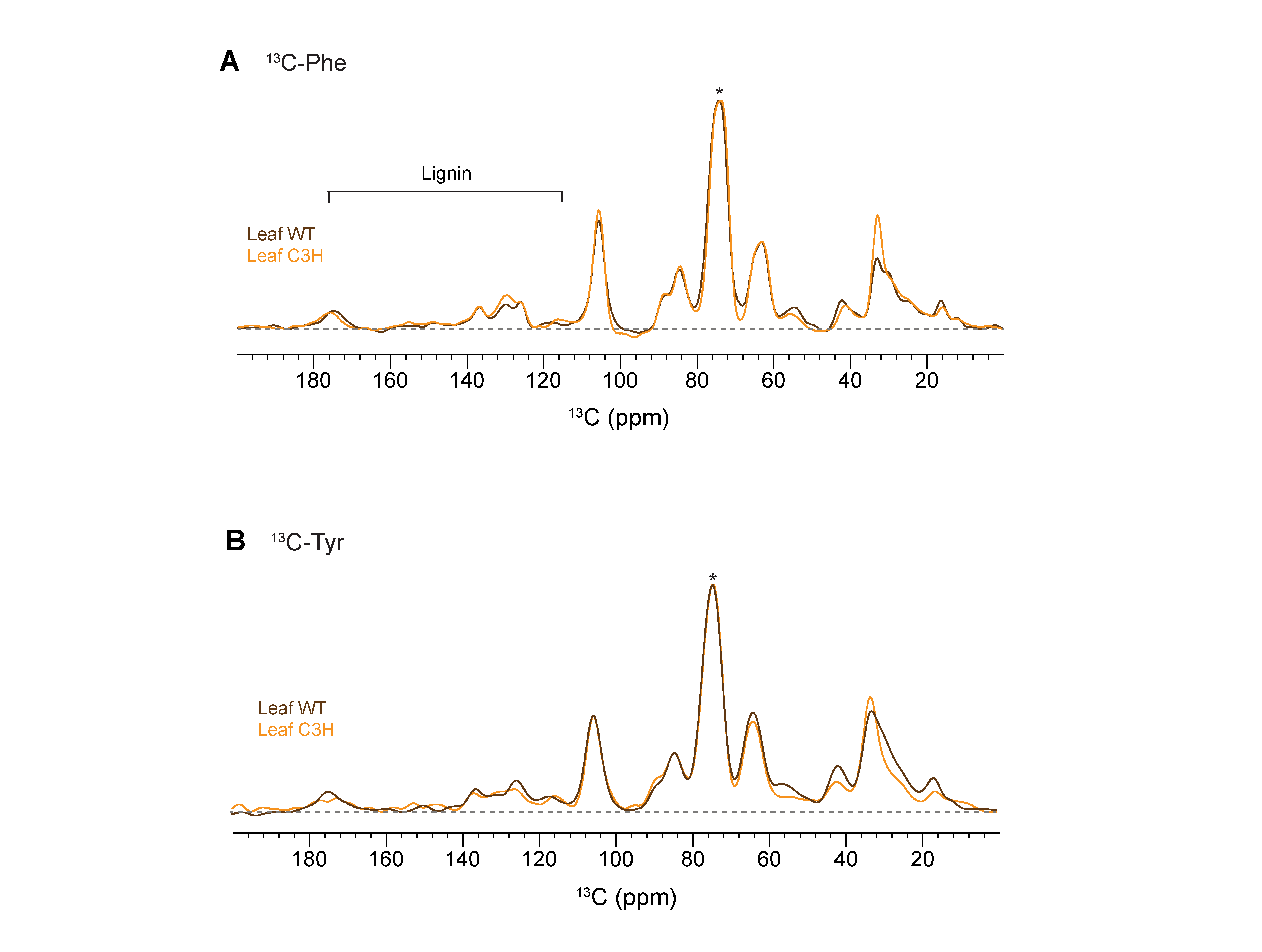

**Figure S6. Leaf tissues show minimal aromatic labeling regardless of genotype.** (**A**) Overlay of 1D ^13^C CP spectra of WT (brown) and C3H (orange) mutant leaf tissues labeled with ^13^C-Phe. (**B**) Corresponding spectra from ^13^C-Tyr labeled leaves. Both sets show extremely low aromatic signals, and no significant difference between WT and C3H mutant. This further supports tissue specificity in lignin incorporation and validates root as the primary site for structural characterization for this study.

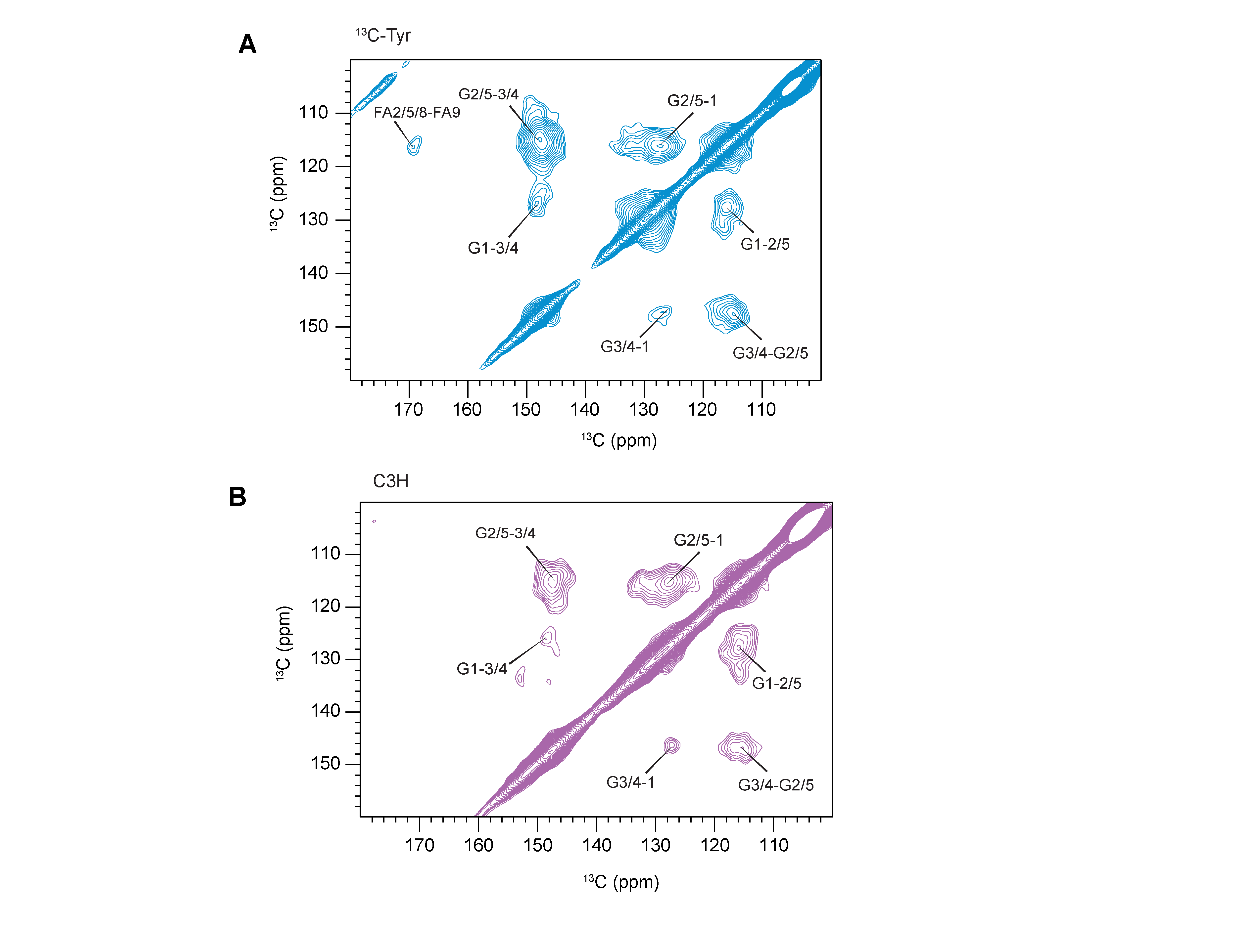

**Figure S7. Medium-contour 2D ^13^C-^13^C solid-state NMR spectra of root tissues labeled with ^13^C-tyrosine** **(A)** WT (blue) and **(B)** C3H mutant (purple). At medium contour levels, both spectra are dominated by G-lignin correlations. A ferulate cross peak at ~169-116 ppm is observed in the WT spectrum but is absent in the C3H mutant, indicating reduced ferulate content in the tyrosine-labeled mutant compared to WT.

**Supplementary Table 1. NMR experimental parameters for all samples.** T = probe temperature; B_0_ = magnetic field; ν_r_ = MAS frequency; NS = number of scans; d_1_ = recycle delay; $\tau$_HC_ = initial ^1^H/^13^C CP contact time; t_2_/t_1_ = acquisition length of direct/indirect dimension; t_m_ = mixing time; TD2/TD1 = total points of FID for the direct/indirect dimension acquisition; TDeff = effective FID data points used for Fourier transform; LB = line broadening parameter; GB = position of the maximum of the Gaussian function; NA = not applicable.

|  | Experiment | T  (K) | B_0_  (T) | *ν*_r_  (kHz) | NS | d_1_  (s) | τ_HC_  (ms) | t_m_  (ms) | t_2_/t_1_  (ms) | TD2/TD1 | DNP Juice | LB  (Hz) | GB |
| --- | --- | --- | --- | --- | --- | --- | --- | --- | --- | --- | --- | --- | --- |
| Conventional  ssNMR (All samples) | 1D CP | 283 | 14.1 | 15.0 | 15k | 2.0 | 1.0 | NA | 15.0 | 1500 | NA | -50 | 0.01 |
| DNP (^13^C-Phe labeled WT) | 1D CP | 110 | 14.1 | 10.5 | 64 | 1.29 | 1.0 | NA | 41.0 | 4096 | d₈-glycerol, D₂O, H₂O (6:3:1) | -50 | 0.01 |
| DNP (^13^C-Tyr labeled WT) | 1D CP | 110 | 14.1 | 10.5 | 64 | 1.29 | 1.0 | NA | 41.0 | 4096 | d₈-glycerol, D₂O, H₂O (6:3:1) | -50 | 0.01 |
| DNP (^13^C-Phe labeled WT) | 2D DARR | 110 | 14.1 | 10.5 | 160 | 1.29 | 1.0 | 50 | 11.2/3.4 | 1024/316 | d₈-glycerol, D₂O, H₂O (6:3:1) | -50/-50 | 0.03/0.03 |
| DNP (^13^C-Tyr labeled WT) | 2D DARR | 110 | 14.1 | 10.5 | 128 | 1.29 | 1.0 | 50 | 11.2/3.4 | 1024/316 | d₈-glycerol, D₂O, H₂O (6:3:1) | -50/-50 | 0.03/0.03 |
| DNP (^13^C-Phe labeled C3H) | 1D CP | 110 | 14.1 | 11.2 | 512 | 1.82 | 1.0 | NA | 41.0 | 4096 | d₈-glycerol, D₂O, H₂O (6:3:1) | -50 | 0.01 |
| DNP (^13^C-Tyr labeled C3H) | 1D CP | 110 | 14.1 | 11.2 | 512 | 1.29 | 1.0 | NA | 41.0 | 4096 | d₈-glycerol, D₂O, H₂O (6:3:1) | -50 | 0.01 |
| DNP (^13^C-Phe labeled C3H) | 2D DARR | 110 | 14.1 | 11.2 | 64 | 1.82 | 1.0 | 50 | 12.3/4.4 | 1200/320 | d₈-glycerol, D₂O, H₂O (6:3:1) | -50/-50 | 0.03/0.03 |
| DNP (^13^C-Tyr labeled C3H) | 2D DARR | 110 | 14.1 | 11.2 | 128 | 1.29 | 1.0 | 50 | 12.3/4.4 | 1200/320 | d₈-glycerol, D₂O, H₂O (6:3:1) | -50/-50 | 0.03/0.03 |

**Supplementary Table 2.** **^13^C chemical shifts of lignin in brachypodium cell walls.** Not applicable (/).

| Lignin | C1 | C2 | | C3 | C4 | C5 | C6 | C7 | C8 | C9 | Experiment | References |
| --- | --- | --- | --- | --- | --- | --- | --- | --- | --- | --- | --- | --- |
| G | 127.4 | | 115.2 | 147.7 | 147.7 | 115.2 | 120.3 | / | / | / | ^13^C-^13^C DARR | Kirui et al. 2022 (1)  Kang et al. 2019 (2)  Xiao et al.  2025 (3) |
| H | 131.3 | | 127.1 | 116.4 | 159.5 | 116.4 | 127.1 | / | / | / |  |  |
| S | 134.4 | | 104.2 | 153.6 | 133.1 | 153.6 | 104.2 | / | / | / |  |  |
| FA | 127.1 | | 116.1 | 148.2 | 148.2 | 116.1 | 127.1 | 148.2 | 116.1 | 168.7 |  |  |

**Supplementary References**

1.	A. Kirui *et al.*, Carbohydrate-aromatic interface and molecular architecture of lignocellulose. *Nat. Commun.* **13**, 538 (2022).

2.	X. Kang *et al.*, Lignin-polysaccharide interactions in plant secondary cell walls revealed by solid-state NMR. *Nat. Commun.* **10**, 347 (2019).

3.	P. Xiao *et al.*, Emergence of lignin-carbohydrate interactions during plant stem maturation visualized by solid-state NMR. *Nat. Commun.* **16**, 8010 (2025).
